# *Burkholderia vietnamiensis* Genes Involved in Extracellular Medium-Chain-Length Polyhydroxyalkanoate Degradation

**DOI:** 10.1101/2025.05.20.655153

**Authors:** Zhong Ling Yap, Sion Yi, Ernesto Quintana, Gabriel G. Lucas, Mohamad Safklou, Ryan Darragh, Andrew M. Hogan, Anna Motnenko, Warren Blunt, Richard Sparling, Dario Fernández Do Porto, David B. Levin, Silvia T. Cardona

## Abstract

Bioplastics represent promising alternatives to petroleum-based plastics, yet their biodegradation remains insufficiently understood. Identifying bacteria capable of degrading bioplastics extracellularly could enhance end-of-life management practices. To investigate *Burkholderia*’s capacities for the degradation of medium-chain-length polyhydroxyalkanoate (mcl-PHA), we screened a panel of *Burkholderia* strains and identified such capacity in strains of *B. gladioli*, *B. multivorans,* and *B. vietnamiensis*. To elucidate the genetic basis of this activity, we performed transposon mutagenesis followed by activity-based screening and Tn-seq on *B. vietnamiensis* LMG 16232. Disrupted genetic elements in transposon mutants with negative phenotypes were further investigated using a CRISPR-associated transposase (CAST) system. These included a lipase production gene cluster, encoding two putative triacylglycerol lipases and a chaperone, and genes coding for a a A24 family peptidase, a TetR/AcrR family transcriptional regulator and a type II secretion system (T2SS) protein. Complete loss or reduced extracellular mcl-PHA depolymerase activity was observed in the CAST mutants, validating their involvement in mcl-PHA degradation. Notably, only one of the two lipases encoded in the lipase production gene cluster was responsible for mcl-PHA degradation, suggesting that while lipases may show substrate promiscuity, lipase functional annotation does not necessarily imply mcl-PHA depolymerization. Docking experiments using the amino acid sequences of the two lipases supported these findings. Together, we identify a gene coding for an active mcl-PHA depolymerase in *B. vietnamiensis* and demonstrate the power of combining activity-based screening, Tn-seq, and CAST to rapidly establish gene-to-function links.

**Importance:** Due to their versatile metabolism, *Burkholderia* strains play critical roles in degradation of multiple compounds in the environment. Here we show that several *Burkholderia* species can extracellularly degrade medium-chain-length polyhydroxyalkanoates (mcl-PHAs), a promising class of bioplastics. By integrating transposon mutagenesis, Tn-seq, and CRISPR-associated transposase (CAST) technologies, we identify and validate key genetic determinants involved in mcl-PHA degradation in *B. vietnamiensis*. These genes encode a lipase, a secretion system component, and regulatory factors, underscoring the complexity and specificity of microbial bioplastic degradation pathways. These findings not only advance our understanding of PHA biodegradation but also identifies *B. vietnamiensis* as as a source of enzymes capable of degrading extracellular mcl-PHA.

## Introduction

Environmental pollution from petroleum-based plastic has driven growing interest in bioplastics, which are bio-based, biodegradable substitutes. Among these, poly(3-hydroxyalkanoates) (PHAs) are microbially produced polyesters that have a wide-range of applications in various sectors, such as packaging, agriculture, and biomedical devices (1). PHAs are synthesized by many microorganisms for intracellular energy and carbon storage under conditions of nutrient imbalance, namely nitrogen and phosphorus limitation. Depending on the number of carbon atoms in each 3-hydroxyalkanoate monomer unit, PHAs can be broadly classified as either short-chain-length PHAs (scl-PHA), which consist of 3–5 carbons per monomer or medium-chain-length PHAs (mcl-PHA), which contain, 6–14 carbons per monomer, each with different physico-chemical properties (2). The carbon side chain length of each monomer will influence the enzyme-substrate specificity for degradation (3).

PHAs are degraded by either intracellular or extracellular PHA depolymerases. Intracellular PHA depolymerases hydrolyze PHA or carbonosomes, which are intracellular carbon reservoirs that can contain PHA granules (4). Extracellular PHA depolymerases degrade exogenous denatured granules that are semi-crystalline and lack the surface layer (3). Both intracellular and extracellular PHA depolymerases are carboxylesterases and belong to the α/β-hydrolase fold family (5). PHA depolymerases have been isolated and biochemically characterized in Firmicutes, Proteobacteria, Actinobacteria, and Ascomycete fungi (6).

Many studies have reported microorganisms that degrade scl-PHA, and have purified and characterized the corresponding enzymes (5, 6). However, microorganisms that can degrade mcl-PHA are relatively rare compared to those that can degrade scl-PHA (6). Most of the extracellular mcl-PHA depolymerases that have been purified and biochemically validated belong to *Pseudomonas* species (7–9), the obligate predator *Bdellovibrio bacteriovorus* (10), *Thermus thermophilus* (11) and *Streptomyces* species (12–14). Amino acid sequence similarity searches of metagenomes using biochemically validated extracellular mcl-PHA depolymerases as queries, have identified candidate extracellular mcl-PHA depolymerases only in the taxonomic groups Myxococcota, Proteobacteria, Actinobacteriota and Bdellovibrionota (6). These studies suggest that mcl-PHA degradation is not widely distributed among Bacteria.

While all extracellular PHA depolymerases hydrolyze PHA, extracellular mcl-PHA depolymerases show little overall sequence homology with scl-PHA. However, PHA depolymerases share structural and mechanistic characteristics with lipases and other α/β-hydrolase enzymes (15). Both PHA depolymerases and lipases have a GxSxG sequence motif found in other α/β-hydrolases and known as the lipase box and a catalytic triad consisting of the central serine of the lipase box, histidine and aspartic acid (16), underscoring a not fully-understood functional promiscuity between PHA depolymerases and lipases. Indeed, some extracellular lipases purified from various species, including *Bacillus subtilis, Pseudomonas aeruginosa, Pseudomonas alcaligenes,* and *Pseudomonas fluorescens* (16) were not able to hydrolyze poly3-hydroxyhexanoate or poly3-hydroxybutyrate. However, the extracellular lipases of *Pseudomonas chlororaphis* PA23 and *Acinetobacter lwoffii* can hydrolyze various PHA polymers, including the mcl-PHA polyhydroxyoctanoate (PHO) and polyhydroxydecanoate (17). While *Burkholderia* strains are known lipase producers (18–21) and the structure of a *B. cepacia* lipase has extensively described (22–26), the capacity to degrade extracellular mcl-PHA in the *Burkholderia* genus has not been investigated.

In this work, we investigated mcl-PHA degradation capacities in *Burkholderia* using PHO as a model substrate. By combining activity-based screening with the power of random and site-directed transposon mutagenesis, we were able to rapidly identify genetic elements that contribute to mcl-PHA degradation in *B. vietnamiensis* LMG 16232. Specifically, we identified the loci P4G95_16805 and P4G95_16810 encoding lipases but only PG95_16805 (*lip1*) is responsible for PHO degradation, while P4G95 1680 (*lip2*) is not. The distinct activities would not have been predicted based on gene annotation alone.

## Results

### Diversity of extracellular mcl-PHA degradation activity in Burkholderia strains

To identify mcl-PHA degradation activity in the *Burkholderia* genus, we performed an initial screen in a solid minimal medium containing PHO and limiting amounts of phenylalanine, supplemented with Nile red to enhance the visualization of PHO degradation as a clear halo around isolated bacterial colonies. Phenylalanine was chosen as a secondary carbon source in the medium because: i) it is known to be catabolized by *Burkholderia* species (27), and ii) its metabolism is known to influence several bacterial pathways (28). The plates were incubated for 20 days to ensure cells grew densely and produced sufficient enzyme to visually detect mcl-PHA degradation. After dense growth was achieved, a clear halo was observed around some colonies, suggesting extracellular PHO degradation.

Several strains of *B. gladioli*, *B. multivorans*, and *B. vietnamiensis,* were able to produce a visible halo suggesting extracellular mcl-PHA degradation (**Table 1, Fig. S1**). While *B. stagnalis* strains MSMB618, MSMB642, and *B. vietnamiensis* LMG 16232 had moderate activity, *B. gladioli* strain VC19233, *B. multivorans* 21NH942533, and VC9825 exhibited weaker activity in degrading mcl-PHA. Two strains, *B. multivorans* F9091687 and *B. gladioli* 132208 observed temperature-dependent mcl-PHA degradation activity. No activity was observed for *B. cenocepacia*, *B. contaminans,* and *B. dolosa* isolates tested (**Table 1**).

**Table 1.**
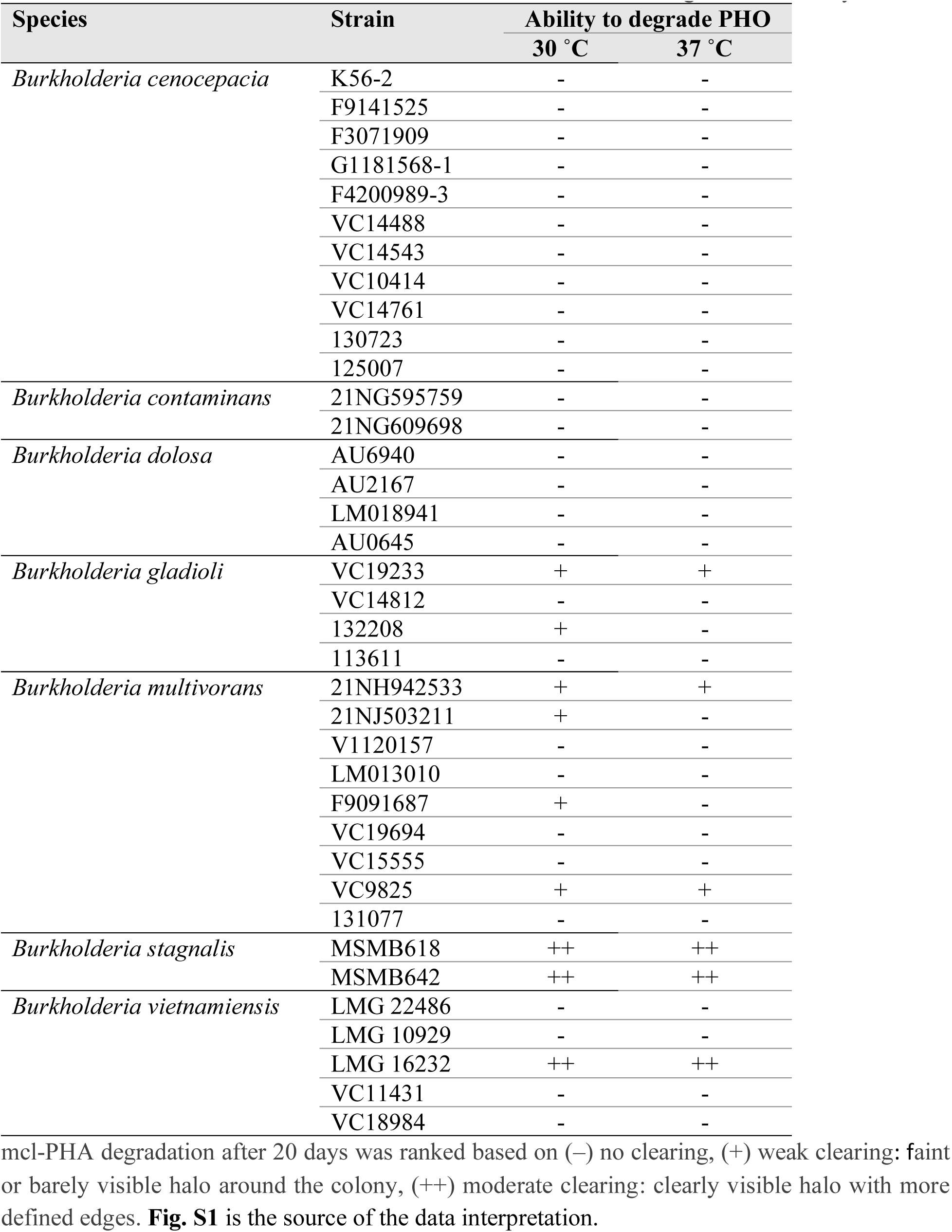
*Burkholderia* isolates that exhibit extracellular mcl-PHO degradation activity.

### Genetic basis of B. vietnamiensis LMG 16232 extracellular mcl-PHA degradation activity

To determine the genetic elements involved in extracellular mcl-PHA degradation activity, we selected *B. vietnamiensis* LMG 16232 (29) for transposon mutagenesis and activity-based screening followed by identification of transposon sites with Illumina sequencing (Tn-seq) (**Fig. 1**). To achieve genomic coverage of transposon insertions in *B. vietnamiensis* LMG 16232, we determined the estimated transposon insertion density using an analytical model based on a Poisson distribution of transposon insertions (30) (see Materials and Methods).

**Fig 1.**
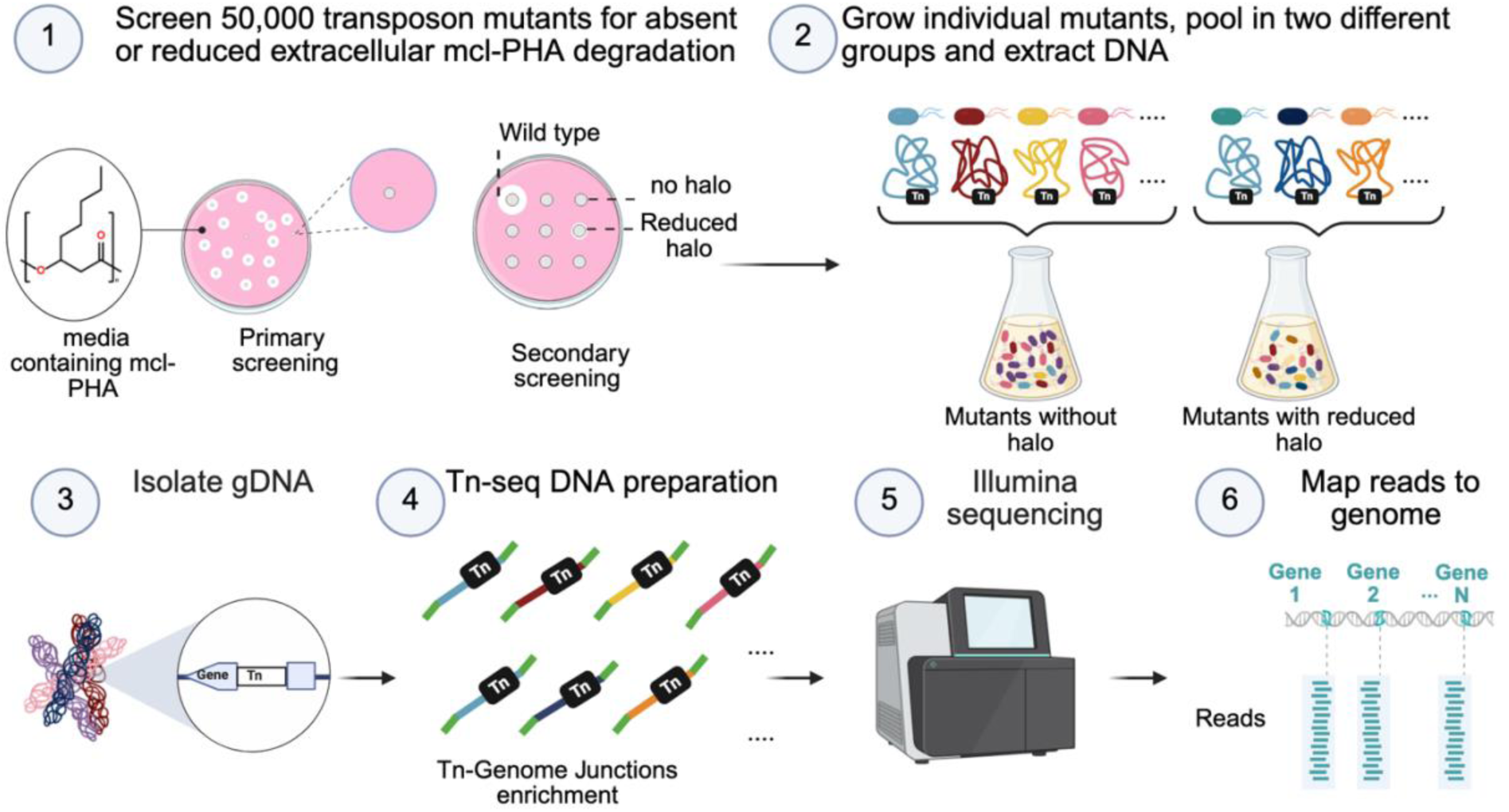
Overview of transposon mutagenesis and Tn-seq workflow to identify genetic elements involved in the extracellular mcl-PHA degradation activity in *B. vietnamiensis* LMG 16232. *B. vietnamiensis* LMG 16232 was subjected to transposon mutagenesis, and 50,000 colonies were screened for those with absent or reduced extracellular mcl-PHA degradation activity. Colonies showing reduced or no degradation activity were selected for further analysis. Selected mutants were grown individually and then grouped into two pools based on their mcl-PHA degradation activity (Pool 1 for no activity, Pool 2 for weak activity). Genomic DNA (gDNA) was extracted from each pool. The gDNA was prepared for Tn-Genome junction enrichment. The enriched DNA fragments were sequenced using Illumina MiSeq. The sequencing data were analyzed by mapping the reads to the *B. vietnamiensis* LMG 16232 genome.

The primary screening of single isolated transconjugants identified 84 colonies that did not produce the visible halo associated with mcl-PHA degradation. These colonies were subjected to secondary screening, where each isolated transconjugant clone was propagated and screened on solid medium containing mcl-PHA. We observed that 66 transconjugants had lost the ability to degrade extracellular mcl-PHA, showing no halo (pool 1). In contrast, 18 transconjugants exhibited a reduced halo (pool 2). Enrichment of transposon insertions followed by Illumina sequencing identified several disrupted genetic elements. Pool 1 and pool 2 contained transposon mutants with insertions in 18 and 12 unique loci, respectively (Table 2).

**Table 2.**
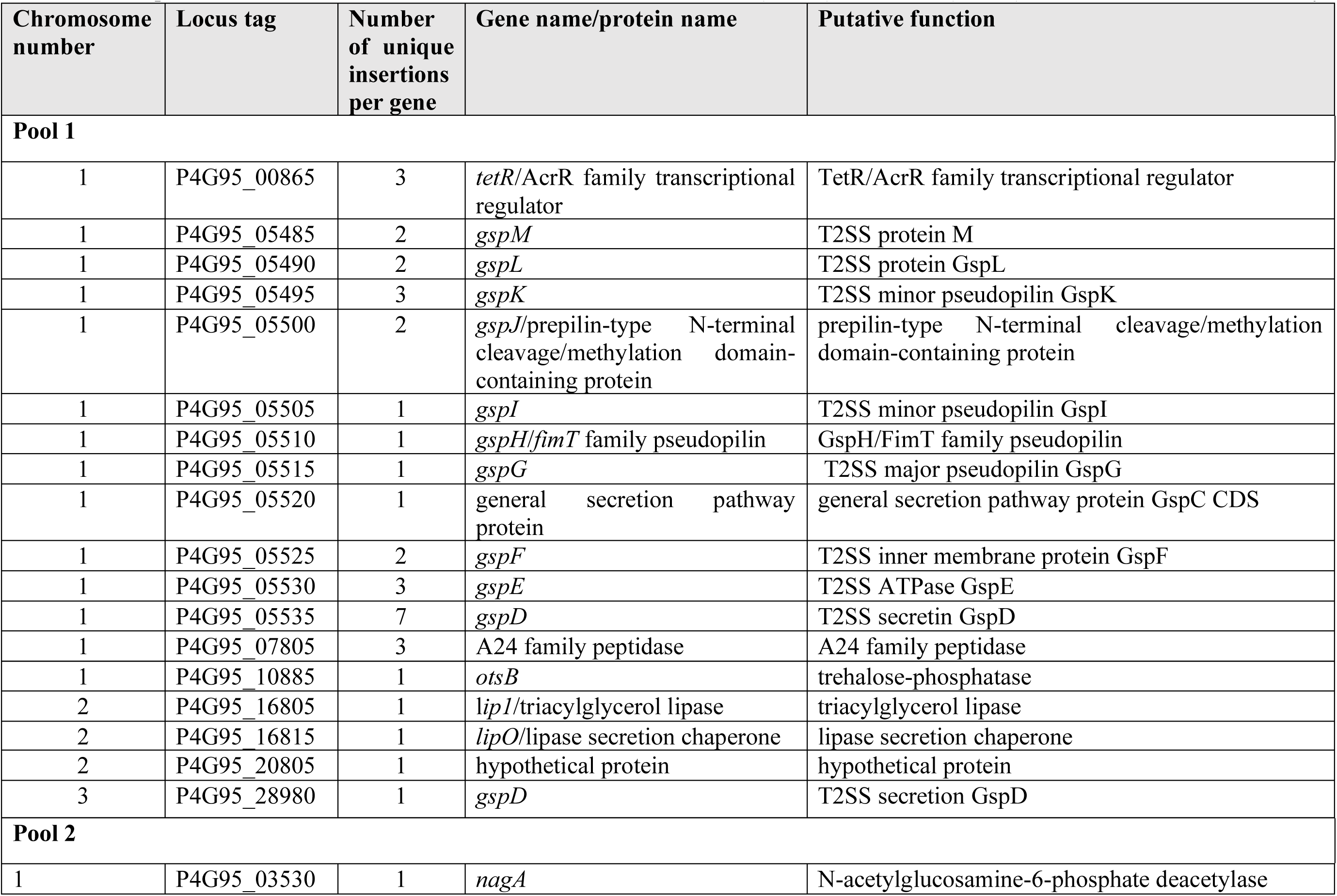
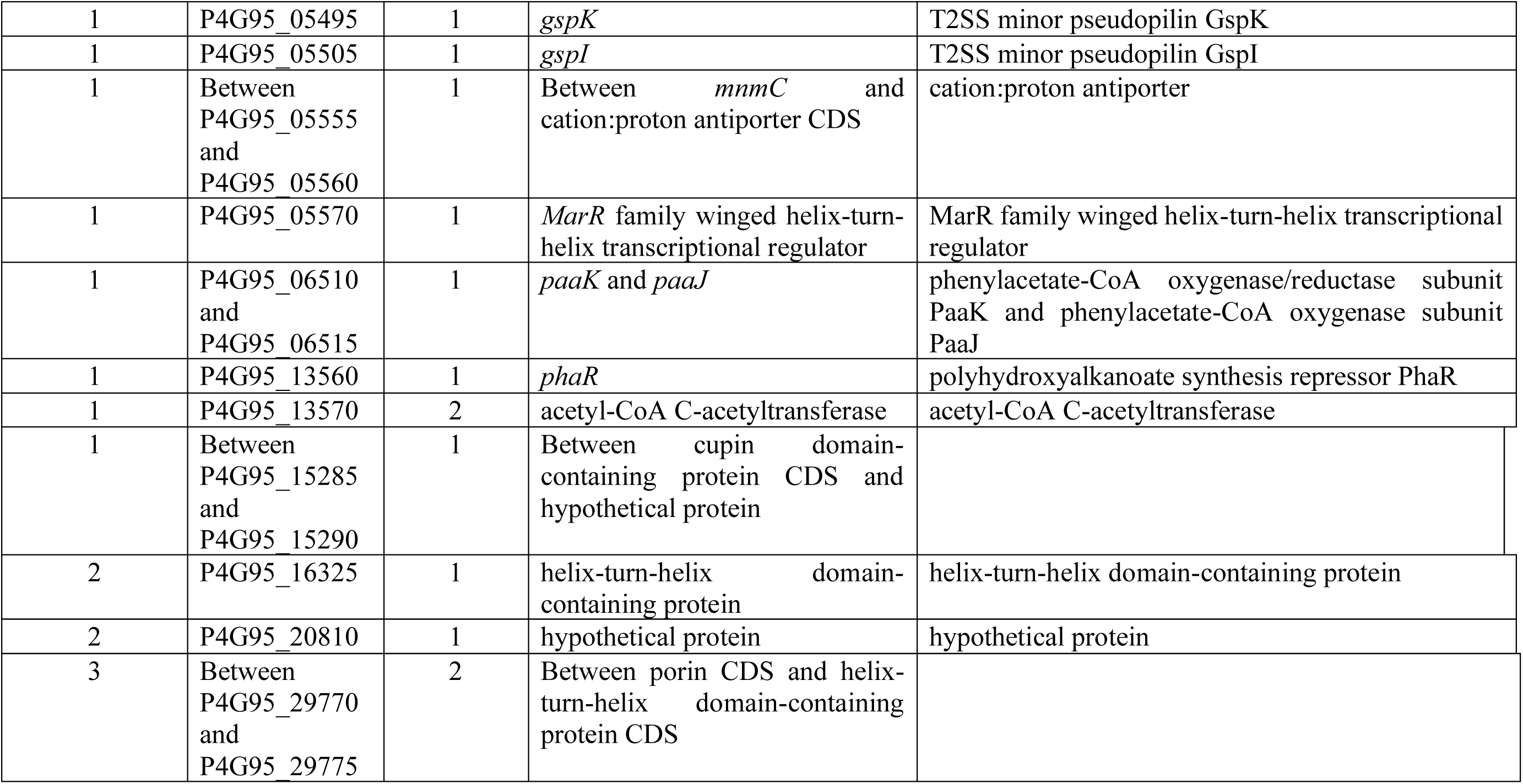
Transposon mutants of *B. vietnamiensis* LMG 16232 with abolished (Pool 1) or reduced (Pool 2) extracellular mcl-PHA activity.

Further examination of the transposon insertions identified gene clusters (P4G95_05485, P4G95_05490, P4G95_05495, P4G95_05500, P4G95_05505, P4G95_05510, P4G95_05515, P4G95_05520, P4G95_05525, P4G95_05530, P4G95_05535) encoding a T2SS (**Fig. 2A**), and a lipase gene cluster, encoding two triacylglycerol lipases and a lipase chaperone (P4G95_16805, P4G16810 and P4G16815, respectively (named *lip1*, *lip2* and l*ipO* herein) (**Fig. 2B**). While activity-abolishing insertions were identified in *lip1* and *lipO*, no transposon insertions that resulted in loss of mcl-PHA degradation were identified in the *lip2* gene, suggesting that *lip1* (P4G95_16805) but not *lip2* (P4G95_16810) might be responsible for extracellular mcl-PHA degradation.

**Fig 2.**
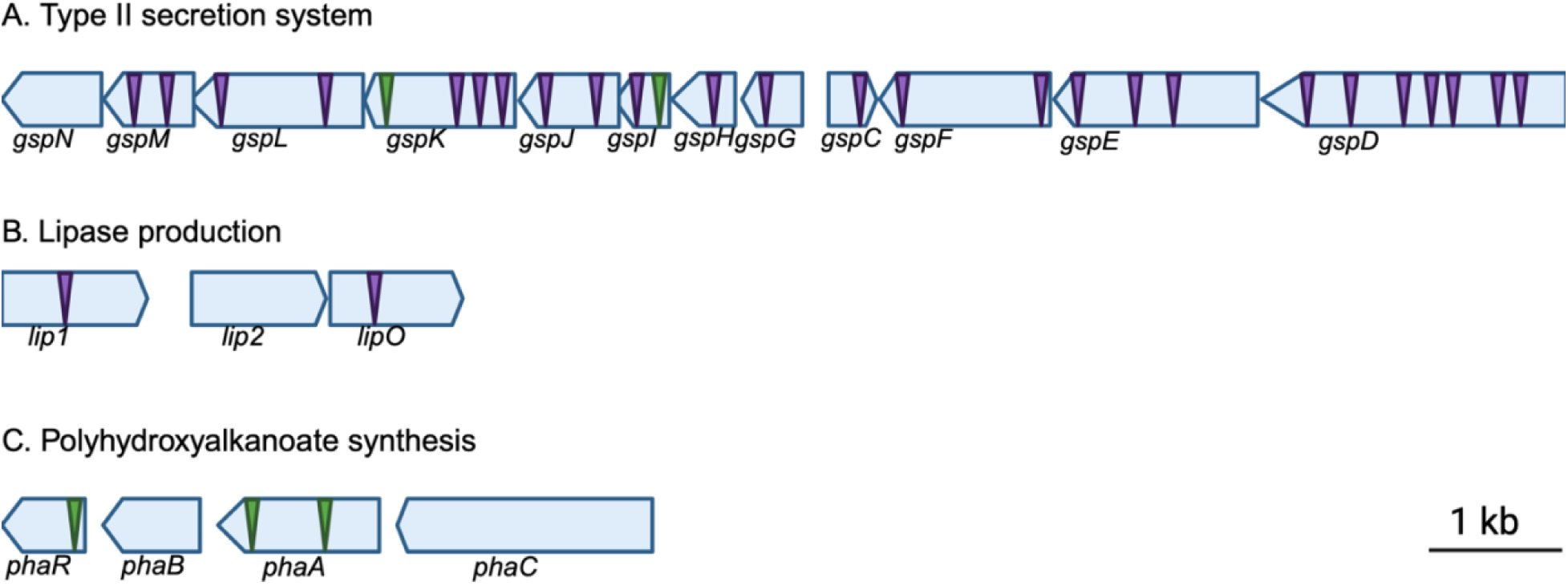
Genetic Organization of the *B. vietnamiensis* LMG 16232 gene clusters that affect the extracellular mcl-PHA when disrupted by transposon insertions. Triangles indicate transposon insertions that resulted in complete loss (purple) or severe reduction (green) of extracellular mcl-PHA degradation activity in *B. vietnamiensis* LMG 16232; **A**) The T2SS of *B. vietnamiensis* LMG 16232 arranges in a gene cluster containing 12 genes (P4G95_05480, P4G95_05485, P4G95_05490, P4G95_05495, P4G95_05500, P4G95_05505, P4G95_05510, P4G95_05515, P4G95_05520, P4G95_05525, P4G95_05530, P4G95_05535). All genes of the gene cluster are interrupted except for *gspN*; **B**) lipase gene cluster consists of three genes, *lip1, lip2* and *lipO* (P4G95_16805, P4G95_16810, and P4G95_16815); **C**) The polyhydroxyalkanoate synthesis gene cluster of *B. vietnamiensis* LMG 16232 consists of four genes, *phaR, phaB, phaA, and phbC* (P4G95_13560, P4G95_13565, P4G95_13570, and P4G95_13575). The genes are depicted to scale, as indicated by the bars on the right side of the panels. The putative function of transposon-disrupted genes is provided in Table 2.

Another identified gene cluster was related to the biosynthesis of polyhydroxybutyrate. Transposon insertion sites in this cluster were found in *phaR* (P4G95_13560), a PHA synthesis repressor and *phaA* (P4G95_13570), which encodes for an acetyl-CoA acetyltransferase involved in the biosynthesis of polyhydroxybutyrate (**Fig. 2C**). In addition, a gene coding for a A24 family peptidase (P4G95_07805) and four genes encoding transcriptional regulators were identified, which are a TetR/AcrR family transcriptional regulator (P4G95_00865), a MarR family winged helix-turn-helix transcriptional regulator CDS (P4G95_05570), *phaR* (P4G95_13560), the polyhydroxyalkanoate synthesis repressor, and a helix-turn-helix domain-containing protein CDS (P4G95_16325) (Table 2). Together, the identified genetic elements suggest the Lip1 lipase, which contains a signal peptide, is cleaved by the A24 family peptidase (P4G95_07805) and is assisted by the LipO lipase chaperone for folding and activation prior to secretion via the T2SS. Finally, the regulatory elements identified together with the *pha* gene cluster suggest that activation of mcl-PHA degradation is regulated and intertwined with PHA biosynthesis.

A few genetic elements obtained from the Tn-seq analysis had a less clear relation with extracellular mcl-PHA degradation in *B. vietnamiensis* LMG 16232. These were genes encoding for trehalose-6-phosphate phosphatase (P4G95_10885), hypothetical proteins (P4G95_20805 and P4G95_20810), the N-acetylglucosamine-6-phosphate deacetylase, NagA (P4G95_03530), a phenylacetate-CoA oxygenase/reductase subunit PaaK and PaaJ (P4G95_06510 and P4G95_06515) and an acetyl-CoA acetyltransferase PhaA (P4G95_13570).

### Genetic elements responsible for mcl-PHA degradation are confirmed by RhaCAST

To validate the genetic elements involved in extracellular mcl-PHA degradation activity identified by Tn-seq, we created independent insertional mutants with our previously developed RhaCAST system (31), which uses a rhamnose-inducible CRISPR-associated transposase (CAST)-based delivery of transposons (32) for targeted insertional mutagenesis in target genes. We applied RhaCAST in *B. vietnamiensis* LMG 16232 (**Fig. 3A**) interrupting genes previously detected by our Tn-seq approach. We successfully produced insertional mutants for genes encoding the putative lipase (*lip1*, P4G95_16805), the A24 family peptidase (*A24*, P4G95_07805), the TetR/AcrR family transcriptional regulator (*tetR/acrR*, P4G95_00865), trehalose-phosphatase (*otsB,* P4G95_10885), and a T2SS protein (*gspE*, P4G95_05530) (**Fig. S2-6**). Inoculation and growth of these mutants on agar plates containing mcl-PHA failed to produce a WT-like visible halo around the colonies (**Fig. 3B**), confirming the link between the interrupted genetic elements and the observed phenotype. To further confirm that the halo of degradation observed corresponds to mcl-PHA degradation, plugs of agar from underneath WT and *lip1* mutant growth were taken and the presence of PHO in the agar was assessed with by gas chromatography with flame ionization detection (GC-FID). As expected, no PHO was detected in the agar beneath the WT strain, whereas PHO was detected under the *lip1* strain, indicating that PHO was not being degraded (**Fig. 4**). We note that after many attempts, we were unsuccessful in producing a RhaCAST *lipO* mutant for unknown reasons.

**Fig 3.**
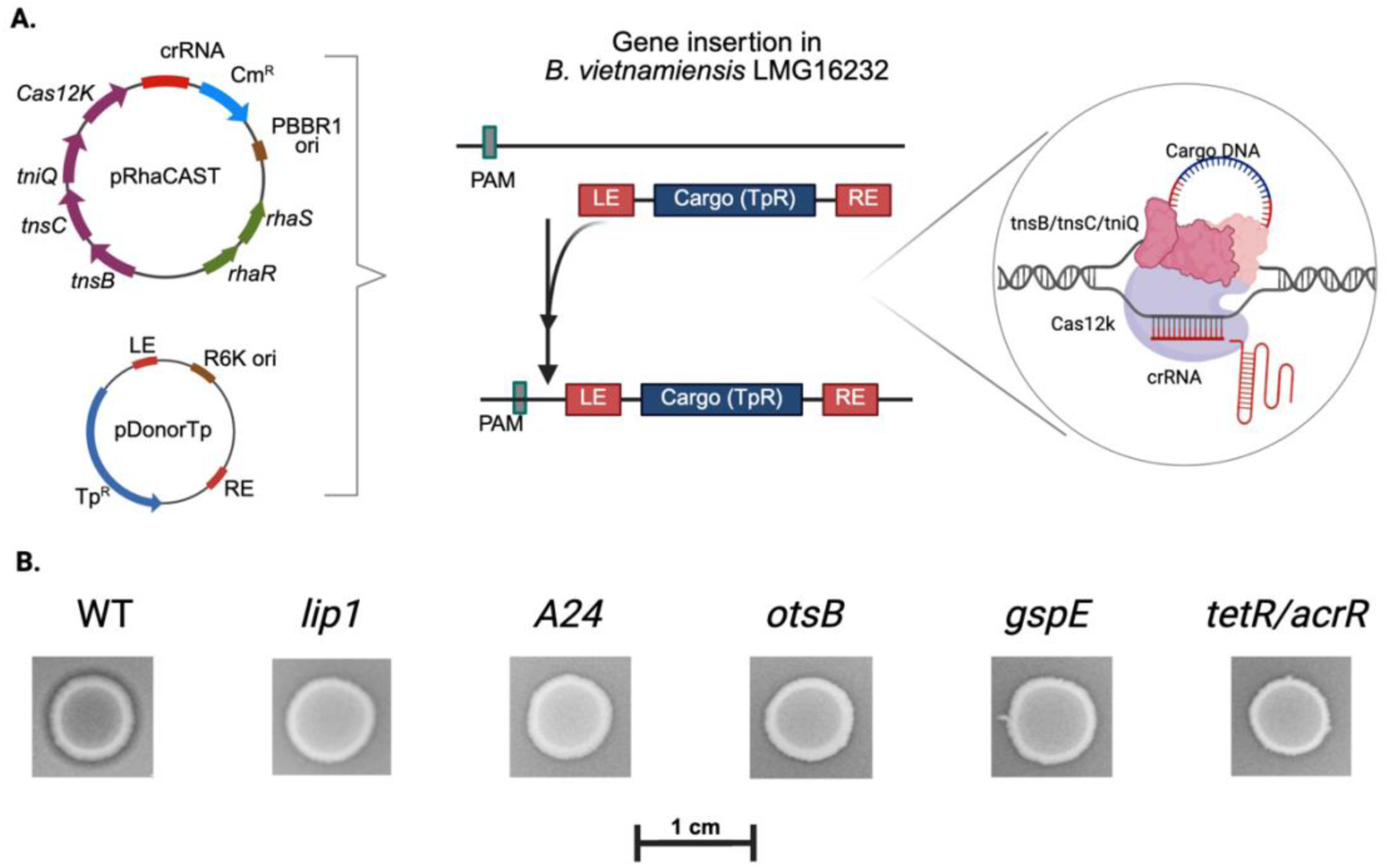
Targeting genes involved in mcl-PHA degradation in *B. vietnamiensis* LMG 16232 abolish the strain’s ability to degrade mcl-PHA; **A**) Schematic model of RNA-guided DNA transposition by RhaCAST applied to *B. vietnamiensis* LMG 16232. The two plasmids of the RhaCAST system, pRhaCAST and pDonorTp, encode all the necessary components, *tnsB*, *tnsC*, *tniQ*, Cas12k, crRNA scaffold, and the cargo gene (TpR) that is flanked by the transposon left end (LE) and right end (RE); **B**) Bacterial growth on PHO plates of WT and RhaCAST mutants. Images are croppings from a representative PHO agar plate. The scale bar (1 cm) is shown for size reference. WT, B. vietnamiensis LMG 16323. *Lip 1,* A24, *otsB, gspE* and *tetR/acrR* are the targeted genes. Putative functions are described in Table 2. Complete name of the mutants is provided in Table 3.

**Fig 4.**
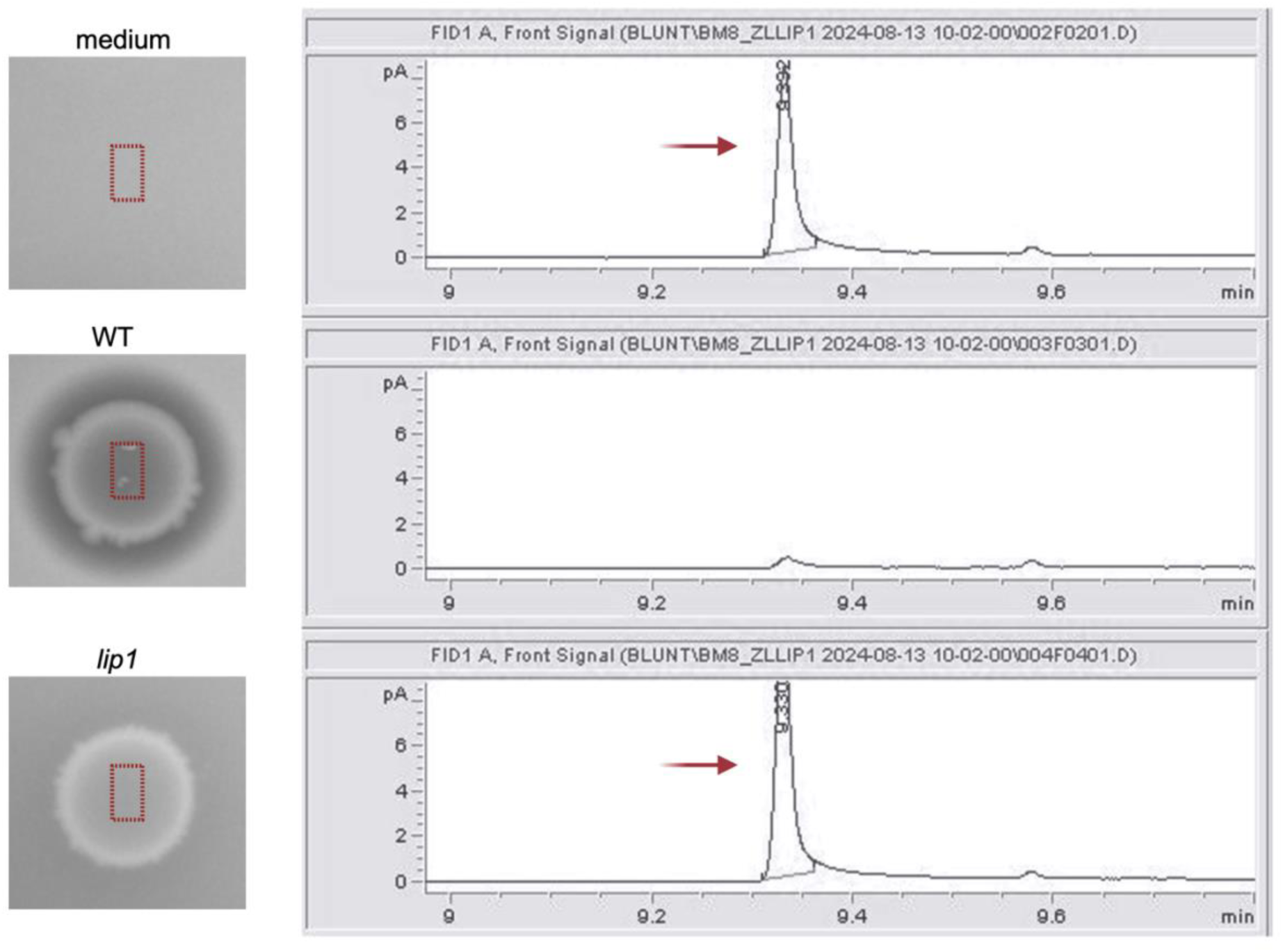
Targeting the putative *lipase* gene (locus tag: P4G95_16805) in *B. vietnamiensis* LMG 16232 abolishes the ability to degrade mcl-PHA. The absence or presence of PHO in the agar after strain inoculation was determined with gas chromatography with flame ionization detection (GC-FID). Samples were obtained by cutting an agar disk under the strain. For control, a disk was cut at an area of the agar where no cell was inoculated. The peak at ∼9.3 minutes retention time corresponds to PHO, which can be observed at the no cell control (Top). The absence of a peak corresponds to PHO in WT, and the presence of PHO underneath the *lip1* mutant growth supports the phenotypic test. Strains were grown on an agar plate containing Nile red and PHO and visualized under UV irradiation. Inoculation of wild type (WT) results in a clear halo around the strain, whereas the inoculation of the *lip1* mutant does not have a halo around the strain.

**Table 3.**
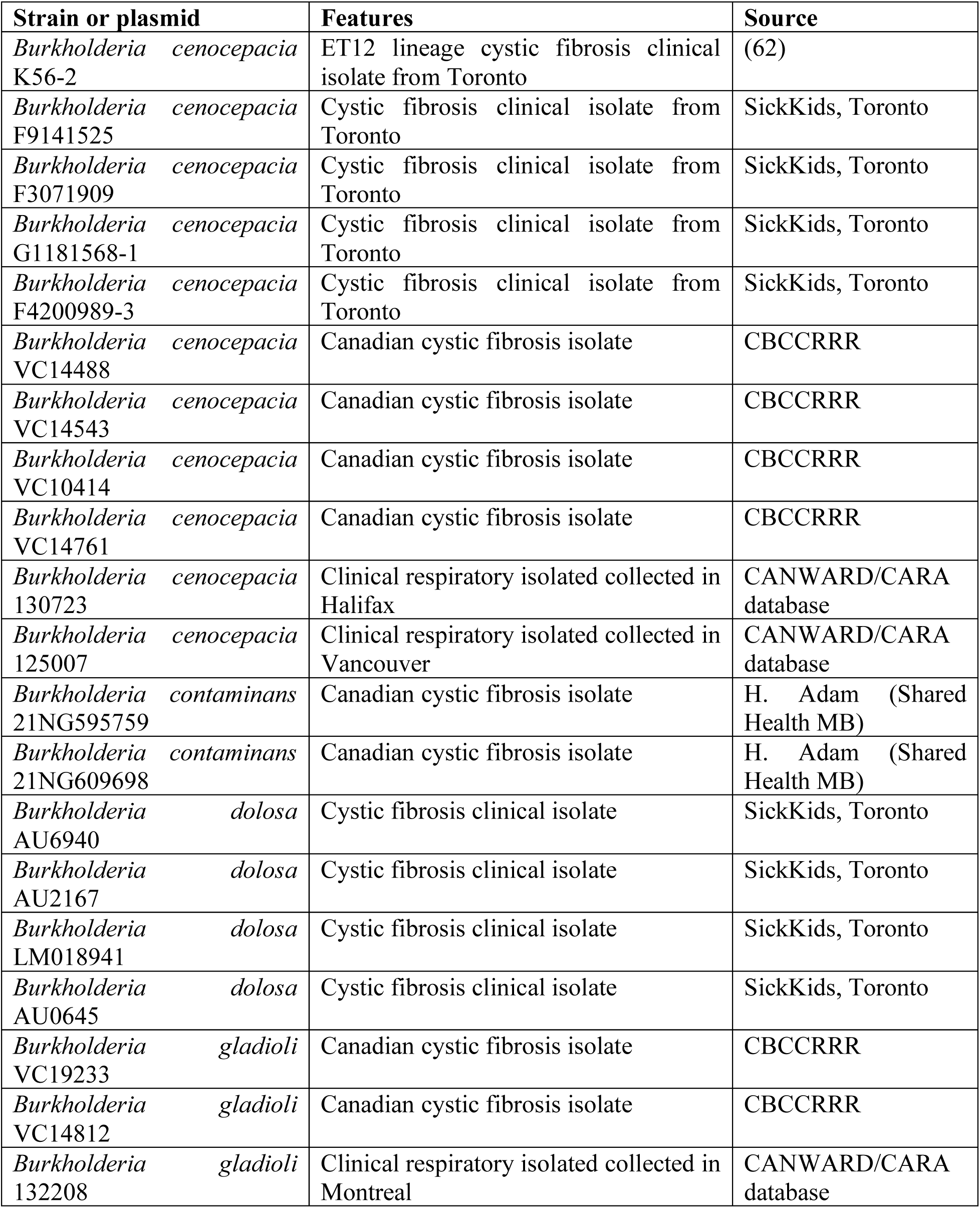
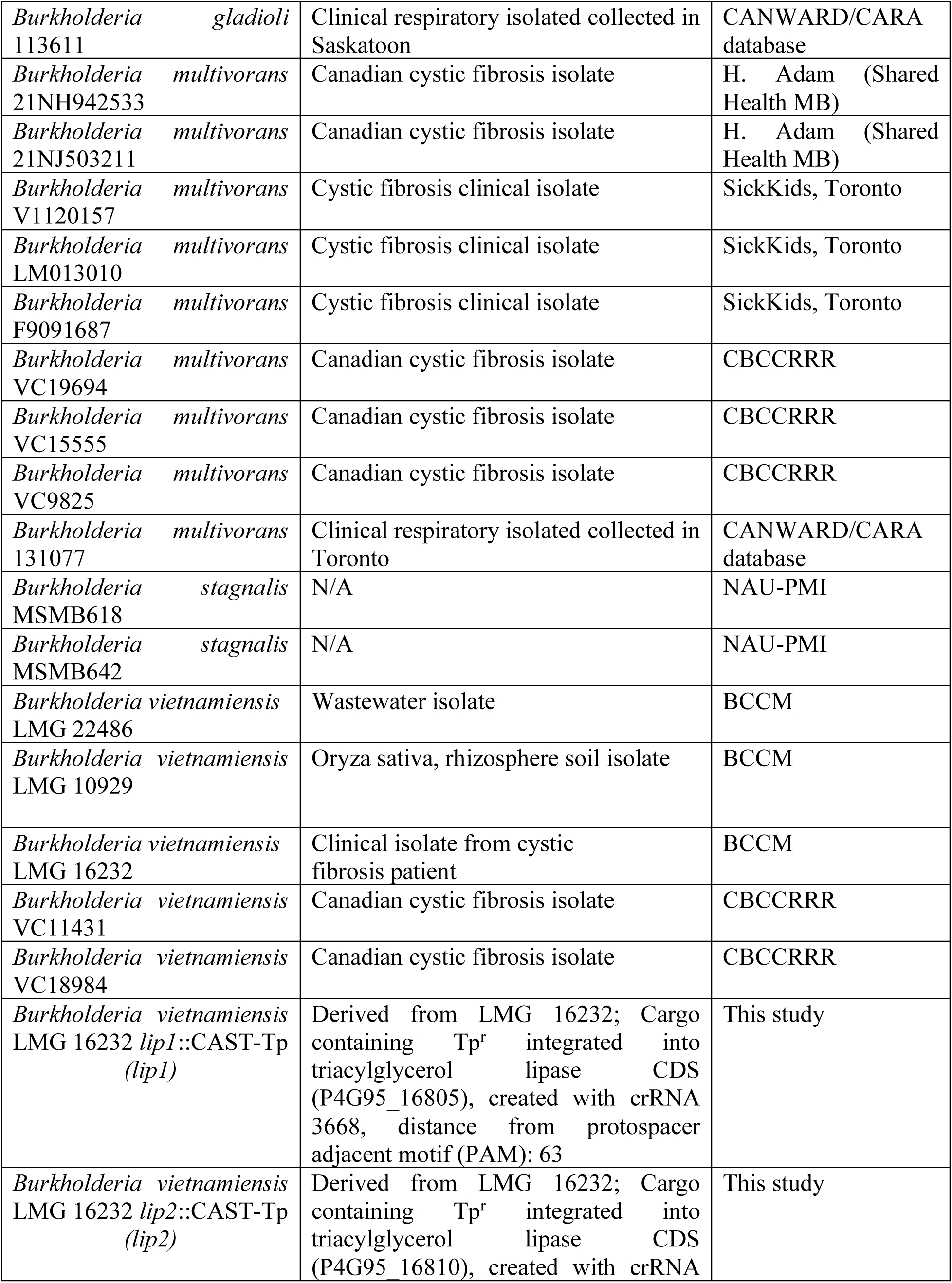
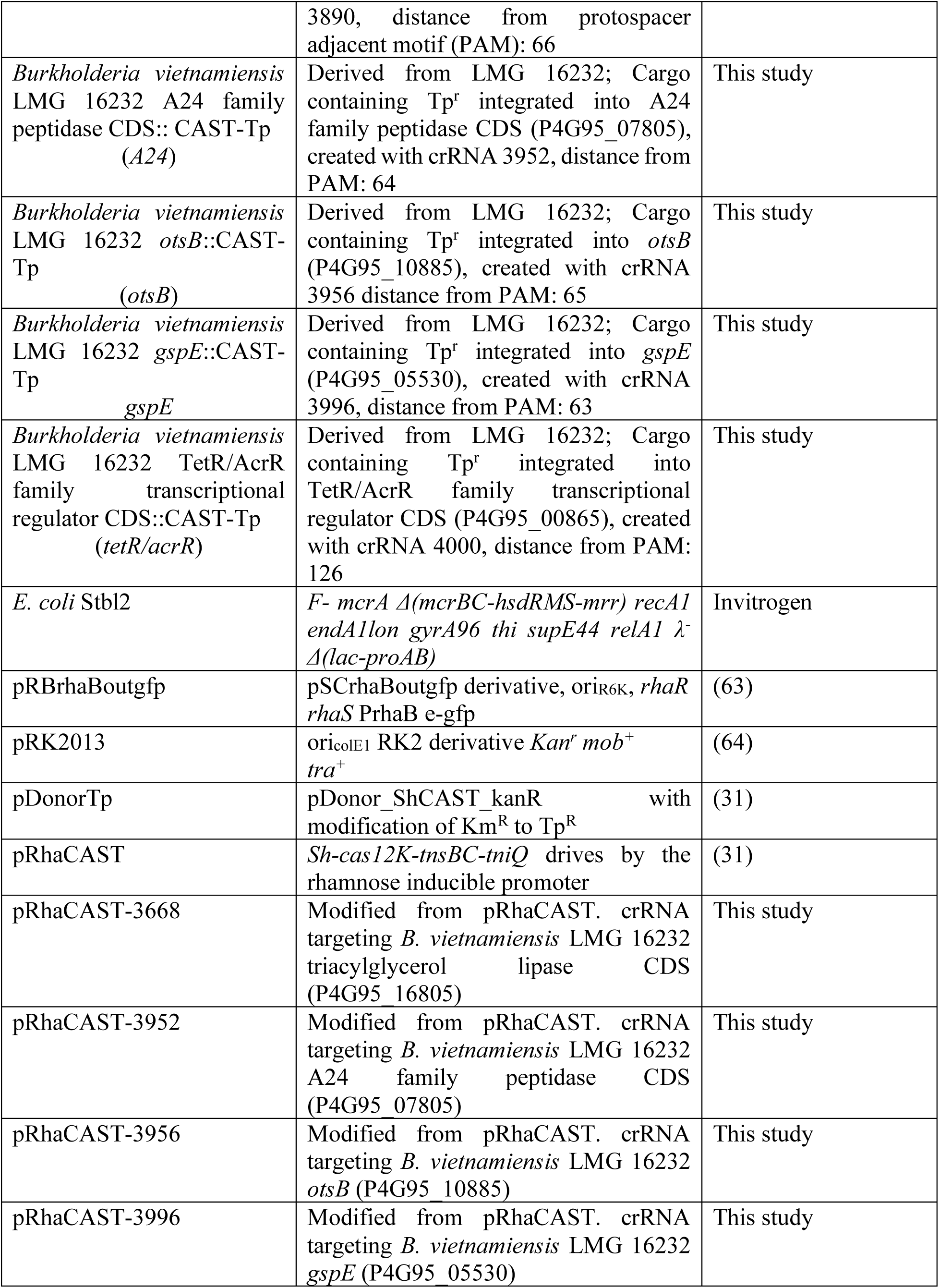
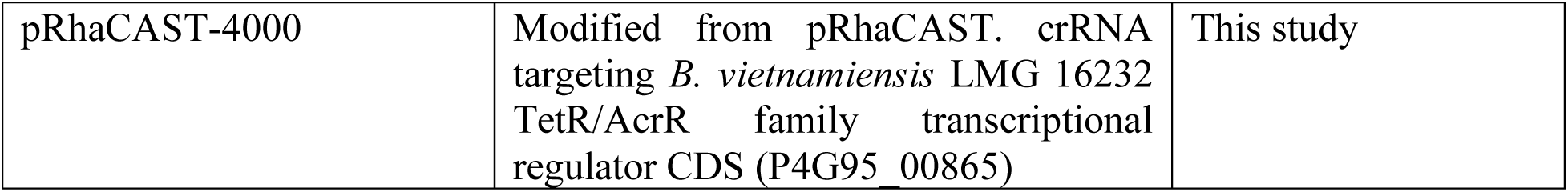
List of bacterial strains and plasmids used in this work.

### Phenotypic and gene expression analysis of the lipase production gene cluster confirms that lip1, but not lip2, is responsible for extracellular mcl-PHA degradation

High-density transposon mutagenesis followed by Tn-seq analysis uncovered the involvement of a lipase production gene cluster (**Fig. 2B** and **Fig. 5A**). From this analysis, colonies with no halo were found to have insertions in *lip1* and *lipO* (**Fig. 5B**) but transposon insertions in *lip2* were not found when sequencing the halo-defective colonies (**Fig. 5B**, N.D). This finding was intriguing for two reasons; first, it seemed that Lip2 had no enzymatic activity against mcl-PHA, despite displaying 78% identity and 86% similarity to Lip1 (**Fig. S7**). Second, as *lip2* and *lipO* seem to be organized in an operon, one might expect that transposon interruptions in *lip2* would result in a halo-defective phenotype because transposon insertions in *lipO* produced no halo around the colonies (**Fig. 5B**). To further elucidate these conflicting results, we used RhaCAST to construct a *lip2* mutant (**Fig. 5B, Fig. S8**). The *lip2* mutant displayed a WT-like halo, indicating mcl-PHA degradation (**Fig. 5B**) and supporting the involvement of only *lip1* in mcl-PHA degradation activity. Furthermore, gene expression analysis of the *lip2* mutant by RT-PCR demonstrated that *lipO* was expressed ruling out a polar effect of the RhaCAST system on this putative operon (**Fig. 5C**). The *lipO* gene was also expressed in the *lip1* mutant (**Fig. 5D**).

**Fig. 5.**
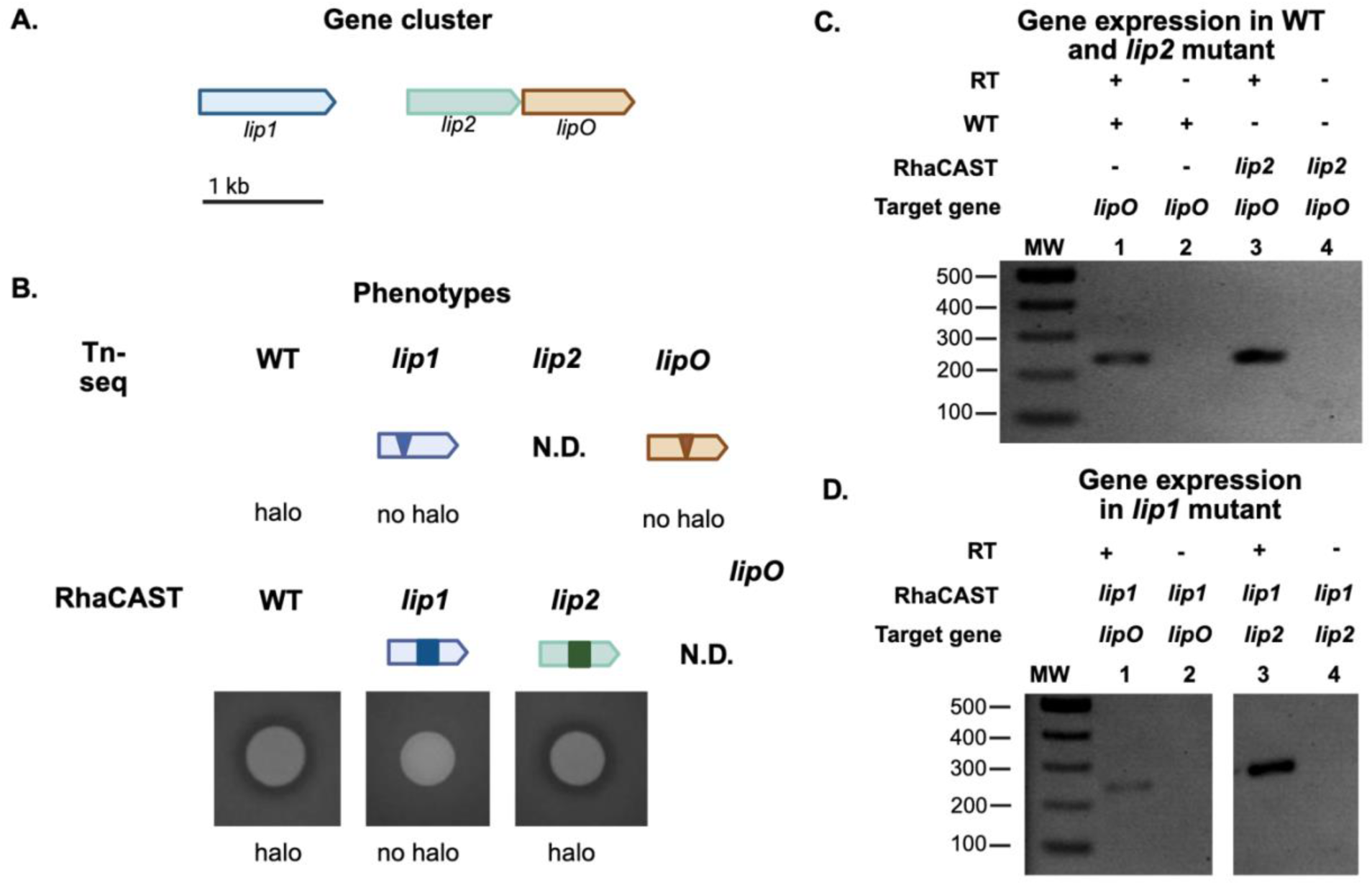
Phenotypic and gene expression analysis of the lipase production gene cluster. **A**) Scheme of the *B. vietnamiensis* LMG 16232 *lip1, lip2* and *lipO* (P4G95_16805, P4G95_16810, and P4G95_16815); **B**) mcl-PHA degradation phenotypes of the respective mutants produced by Tn-seq (scheme) or RhaCAST (bacterial growth on PHO plates after 7 days); **C-D)** Lipase cluster expression analysis. WT, *lip1* and *lip2* mutants were grown on PHO plates and at day 0 (shown) 4 and 7, cells were harvested for RNA extraction. Expression of target genes was confirmed by RT-PCR. RT, reverse transcriptase.

As Lip2 was not involved in extracellular mcl-PHA degradation, we compared the activity of *lip1* and *lip2* mutants towards tributyrin, a well-known substrate for lipase activity (33) and quantified degradation halos of biological replicates (**Fig. 6**). The wild-type *B. vietnamiensis* LMG 16232 exhibited consistent activity against mcl-PHA (**Fig. 6A**). Instead, the *lip1* mutant exhibited no activity against mcl-PHA, confirming its role as solely responsible for mcl-PHA degradation. *B. vietnamiensis* LMG 16232 WT also degraded tributyrin (**Fig. 6B**). However, the tributyrin degrading activity of the *lip1* mutant was only reduced but not abolished (**Fig. 6B**), suggesting that other lipases, including *lip2,* might be active lipases. mcl-PHA and tributyrin degradation were also not affected in the *lip2* mutant (**Fig. 6A and B**). In summary, while Lip1 appears to be active against extracellular mcl-PHA and tributyrin, Lip2 may be active against tributyrin but not against mcl-PHA.

**Fig 6.**
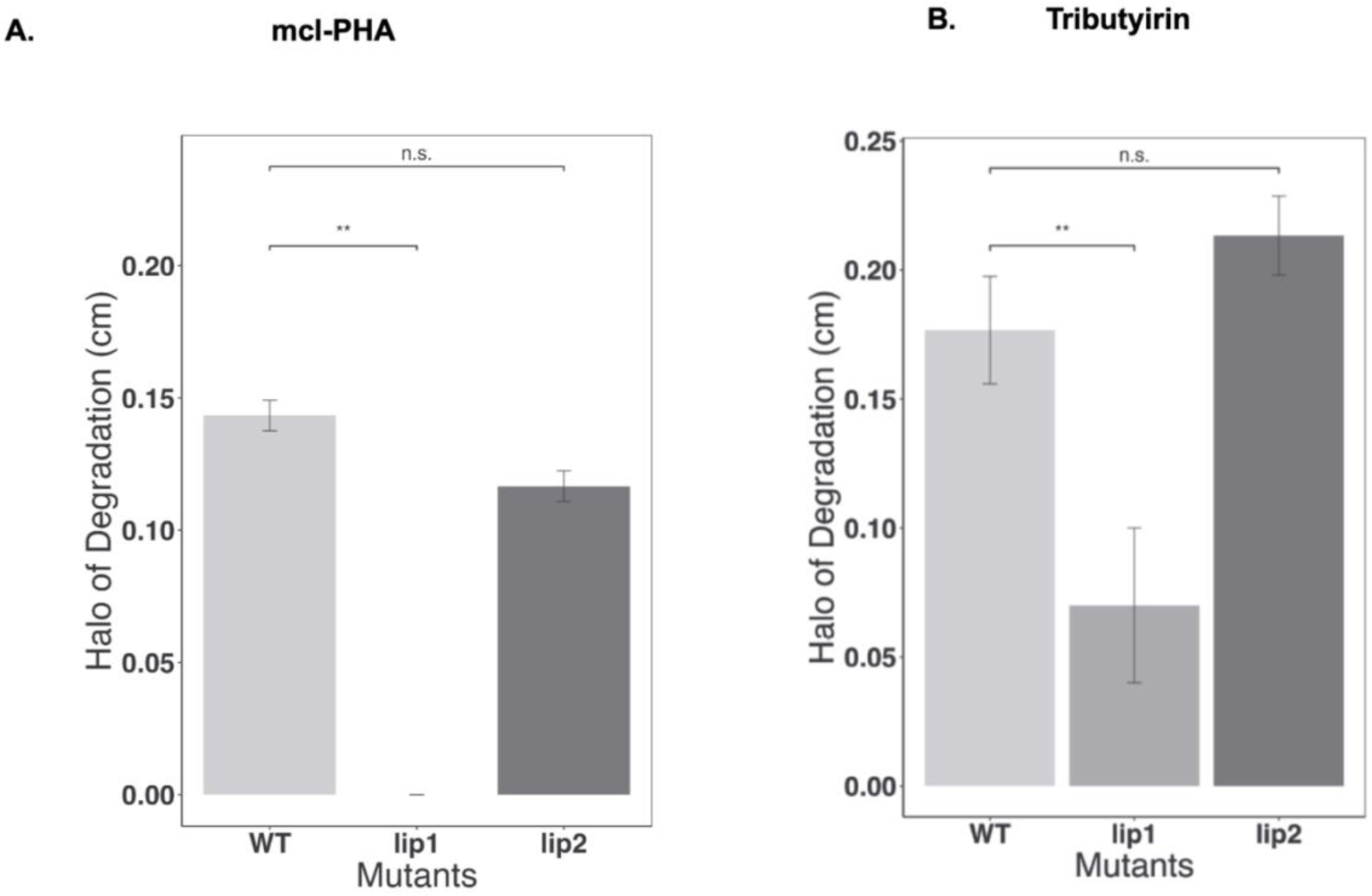
Quantification of mcl-PHA and tributyrin degradation in *B. vietnamiensis* LMG 16232, *lip1* and *lip2* mutants. Diameter of the halos degradation in **A**) mcl-PHA or **B**) tributyrin. Strains were grown on an agar plate supplemented with tributyrin for 3 days or mcl-PHA for 7 days at 37°C. Data is representative of three biological replicates repeated in technical duplicates.

### Structural analysis of Lip1 and Lip2 putative amino acid sequences reveal differences in the catalytic site

To further investigate the differential activity of Lip1 and Lip2, we first compared their sequences with the well characterized *B. cepacia* lipase (22, 24) and experimentally validated mcl-PHA depolymerases from *Streptomyces roseolus* and *S. venezuelae*, (13, 14) and *Pseudomonas* (7–9), as well as *Bdellovibrio* (10). The multiple alignment (**Fig. S9**) revealed that Lip1 and Lip2 showed high similarity to the *B. cepacia* lipase (97.8% and 93.9%, respectively). Instead, less than 30% similarity was found between Lip1 or Lip2 to any of the other mcl-PHA depolymerases beyond the presence of the conserved lipase box. We then predicted the structures of Lip1 and Lip2 using AlphaFold 2 and assessed multiple sequence alignments. The sequence coverage analysis, which combines alignment depth and sequence identity, illustrates the extent of conservation across homologous sequences for both proteins (**Fig. S10A**). A well-conserved core region (residues ∼50–160) is evident in both cases, with lower coverage and diversity in the N- and C-terminal regions suggesting flexible or functionally variable domains. Consistent patterns in Predicted Aligned Error (PAE) matrices of the five top-ranked models support a robust domain architecture with potential inter-domain flexibility (**Fig. S10B**). Additionally, the high predicted Local Distance Difference Test (pLDDT) scores (**Fig. S10C**) further validate the confidence in structural predictions. The high pLDDT values (>90) in the core region confirm a well-defined fold, whereas significant drops around residues 50–100 and 200–250 suggest intrinsically disordered or flexible regions. These trends are consistent across both Lip1 and Lip2, reinforcing the reliability of the predictions and indicating a shared structural organization.

Superposition of the AlphaFold-predicted Lip1 structure (**Fig. S11**) with the *B. cepacia* lipase (PDB ID 4LIP) crystallographic template revealed a high degree of structural similarity except for an alpha-helix, which appears to obstruct a cavity present in 4LIP. To evaluate the catalytic sites in the context of the observed *lip1* and *lip2* mutant phenotypes, we performed a structural superposition with the *B. cepacia* lipase. The overlay reveals a remarkable similarity in the spatial arrangement of the catalytic triad (**Fig. S12**). The conserved positioning of the key catalytic residues strongly suggests that Lip1 and Lip2 adopt a functional fold compatible with lipase activity. This structural conservation provides additional support for the hypothesis that both proteins share mechanistic features with known lipases.

Using the *B. cepacia* lipase as a template, we guided the docking of Lip1 and Lip2 with two substrates: 3-Hydroxyoctanoate (OCT) and Tributyrin (TRI). Both ligands were energy-minimized before docking, and a grid-based docking algorithm was used to position the ligands within the predicted active sites of both proteins. Notably, all combinations of protein-ligand docking yielded clusters with low energy, indicating favourable interactions (**Table S1**).

Docking simulations with Lip1 revealed a single predominant low-energy cluster for both OCT and TRI (**Fig. 7A, B, E, Table S1**). This convergence, together with the consistently low binding energies, strongly suggests a high affinity between Lip1 and both ligands. In contrast, the docking of both ligands against Lip2 (**Fig. 7C, D, E, Table S1**) yielded three distinct low-energy clusters for each substrate. This multiplicity of binding modes suggests a weaker or less specific interaction, potentially due to a more flexible or heterogeneous binding pocket that compromises optimal substrate accommodation. Structural analysis of the catalytic site revealed that Lip1 and the *B. cepacia* lipase share a similar configuration, whereas Lip2 contains a different amino acid at position 73 compared to Lip1, which is glutamine (Lip2) instead of tyrosine (Lip1) (**Fig. S7, Fig. 7F**). This amino acid difference may explain the different phenotypes of the *lip1* and *lip2* mutants in which activity against mcl-PHO was only observed in the *lip2* (but not l*ip1*) mutant.

**Fig. 7.**
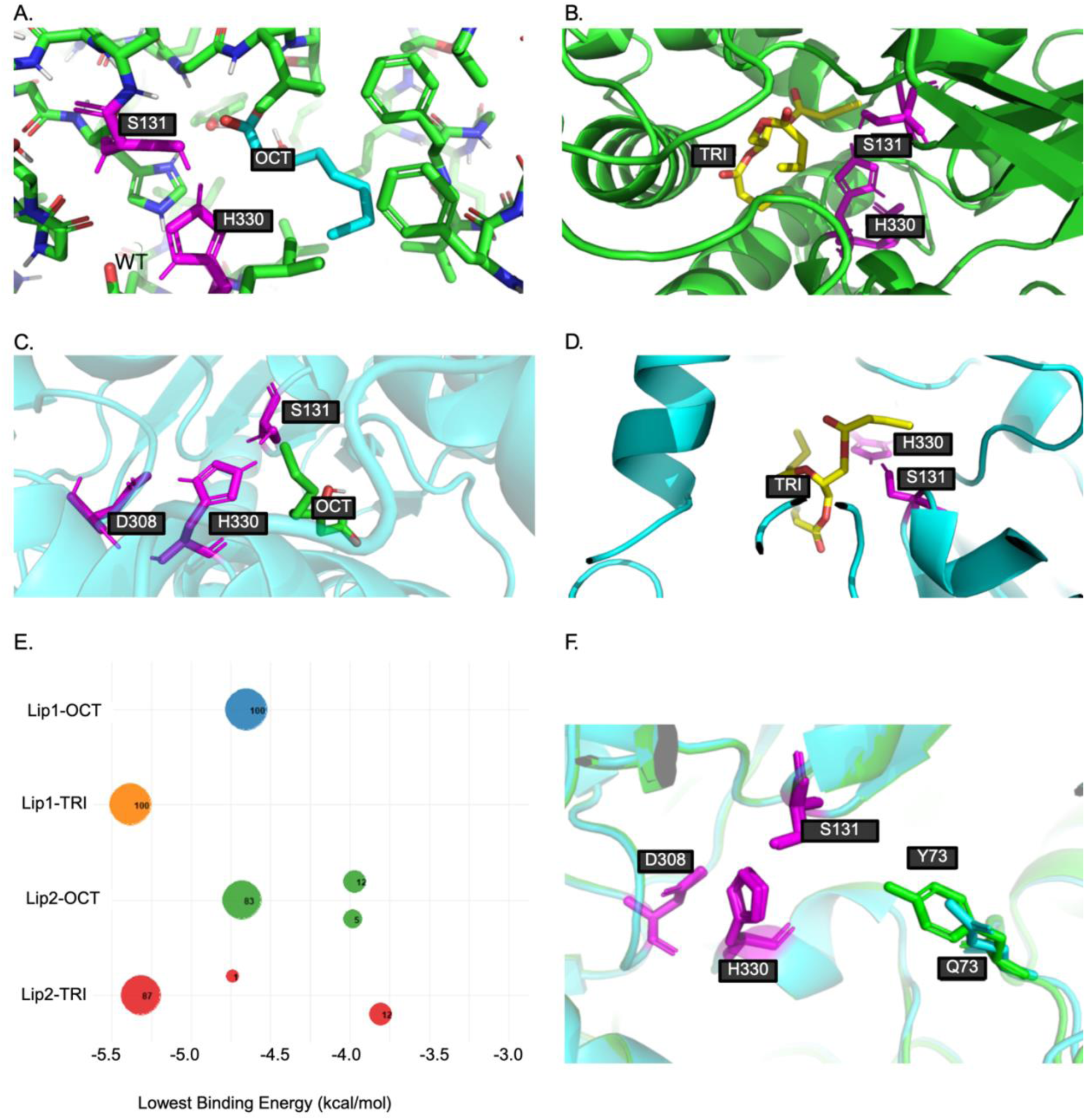
Docking analysis of Lip1 and Lip2-substrate complexes. **A**) Docking poses of Lip1 with 3-Hydroxyoctanoate (OCT); **B**) Docking poses of Lip1 with Tributyrin (TRI); **C**) Docking poses of Lip2 with 3-Hydroxyoctanoate (OCT); **D**) Docking poses of Lip2 with Tributyrin (TRI). The images illustrate the single predominant low-energy binding cluster obtained for both substrates, highlighting the catalytic triad in violet; **E**) Binding energy and cluster distribution for protein-ligand complexes. The plot shows the lowest binding energy for Lip1 and Lip2 docking with 3-Hydroxyoctanoate (OCT) and Tributyrin (TRI). The size of the circle represents the cluster size (number of poses), while colours differentiate protein-ligand pairs: Lip1-OCT (blue), Lip1-TRI (orange), Lip2-OCT (green), and Lip2-TRI (red). Lip1 exhibits a higher affinity for both ligands, characterized by a single predominant low-energy cluster, whereas Lip2 displays multiple binding modes for both substrates; **F**) Structural superposition of Lip1 and Lip2 at amino acid at position 73. Superposition of Lip1 (green) and Lip2 (blue), highlighting the catalytic triad (violet) and the difference in amino acid in position 73: Lip1 (tyrosine, Y) and Lip2 (glutamine, Q), illustrating its potential impact on enzyme structure and function.

## Discussion

The production and use of bioplastics, including mcl-PHA, have increased recently. These materials are considered alternatives to petroleum-based plastics because they can be derived from renewable resources and have a wide range of applications (1). Despite growing interest, microorganisms and enzymes capable of degrading mcl-PHA remain relatively understudied (6). Extracellular mcl-PHA depolymerases have been found in some genera, including *Pseudomonas* (7–9), *Bdellovibrio* (10), *Thermus* (11) and *Streptomyces* (12–14). However, to date, the mechanism of action for extracellular mcl-PHA by microorganisms has not been thoroughly explored. Here, we report that several *Burkholderia* strains, including *B. gladioli*, *B. multivorans*, and *B. vietnamiensis,* can degrade mcl-PHA. Through transposon mutagenesis combined with Tn-seq analysis, we identified genetic elements associated with mcl-PHA degradation in *B. vietnamiensis* LMG 16232. We further validated some of these genetic elements by generating targeted insertional mutants using RhaCAST and tested their extracellular mcl-PHA degradation activity. Our findings indicate that the enzyme responsible for extracellular mcl-PHA degradation is a lipase (Lip1) with substrate versatility and we ruled out the involvement of a second lipase (Lip2) encoded downstream of *lip1*. Indeed, the *lip2* mutant constructed with our RhaCAST system, had similar extracellular mcl-PHA activity compared to the wild type. Therefore, we show experimental evidence of *Burkholderia*’s capability to degrade extracellular mcl-PHA, linking the enzymatic activity to the P4G95_16805 locus (*lip1*) of *B. vietnamiensis* LMG 16232, and ruling out the involvement of *lip2* (P4G95_16810).

A technique commonly used to enhance the visualization of PHA degradation involves incorporating Nile red, a lipophilic fluorescent dye, into agar plates (34). Thus, we adopted this visualization method in our screening medium to increase sensitivity. In comparison to other studies, where the degradation of extracellular mcl-PHA can be observed within 2 to 3 days, *Burkholderia vietnamiensis* LMG16232 exhibited a slower degradation process, requiring approximately 7 days for observable activity, suggesting co-metabolism, and/or poor growth on mcl-PHA. Some strains of *Burkholderia* from the initial screening showed weak activity, with degradation taking up to 20 days, despite the use of Nile red to enhance detection. Nevertheless, by implementing this visualization technique alongside transposon mutagenesis in *B. vietnamiensis* LMG 16232, we were able to screen 50,000 individual colonies in a rapid manner. This approach enabled us to identify transconjugant colonies that had lost the ability to degrade mcl-PHA, providing a sensitive and efficient screening method. This method may be applicable to other polymers that can be stained with suitable dyes, enhancing assay sensitivity.

Besides the identification of *lip1* as the gene encoding an enzyme capable of mcl-PHA degradation, transposon mutagenesis combined with Tn-seq identified other genetic elements involved in the degradation of mcl-PHA in *B. vietnamiensis* LMG 16232 (**Fig. 8**). These genes can be classified into three functional groups: a) lipase/chaperone gene cluster, b) components of T2SS, and c) regulatory elements. The genetic elements associated with the mcl-PHA degradation in *B. vietnamiensis* LMG 16232 are consistent with the findings in various *Burkholderia* lipases, particularly regarding the production, chaperone-assisted folding and secretion of extracellular lipases by T2SSs (35, 36). Similar to the lipases from other *Burkholderia* species, the identified *B. vietnamiensis* LMG 16232 LipO is expected to assist the correct folding and efficient secretion of Lip1 via the T2SS. Many *Burkholderia* strains can secrete extracellular lipases under inducing conditions, and some of those lipases are capable of degrading various compounds (37–40). For example, lipases from *Burkholderia* stains MC 16-3 (GenBank: AY772173) and 99-2-1 (GenBank: AY772174) can degrade methyl(R)-N-(2,6-dimethyl-phenyl)alaninate (41). Additionally, the lipase from *B. cepacia* PBSA-1 (GenBank: EF189714.1) has been shown to degrade poly(butylene succinate-co-butylene adipate) (42), while *B. cepacia* lipase can degrade polycaprolactone (PCL) (16). These lipases, including the ones identified in this work, share the conserved lipase box of G-H-S-Q-G. While *B. cepacia* DP1 is a major lipase producer and has been shown to degrade extracellular scl-PHA (43, 44), there was no report of *Burkholderia* species that degrade extracellular mcl-PHA.

**Fig 8.**
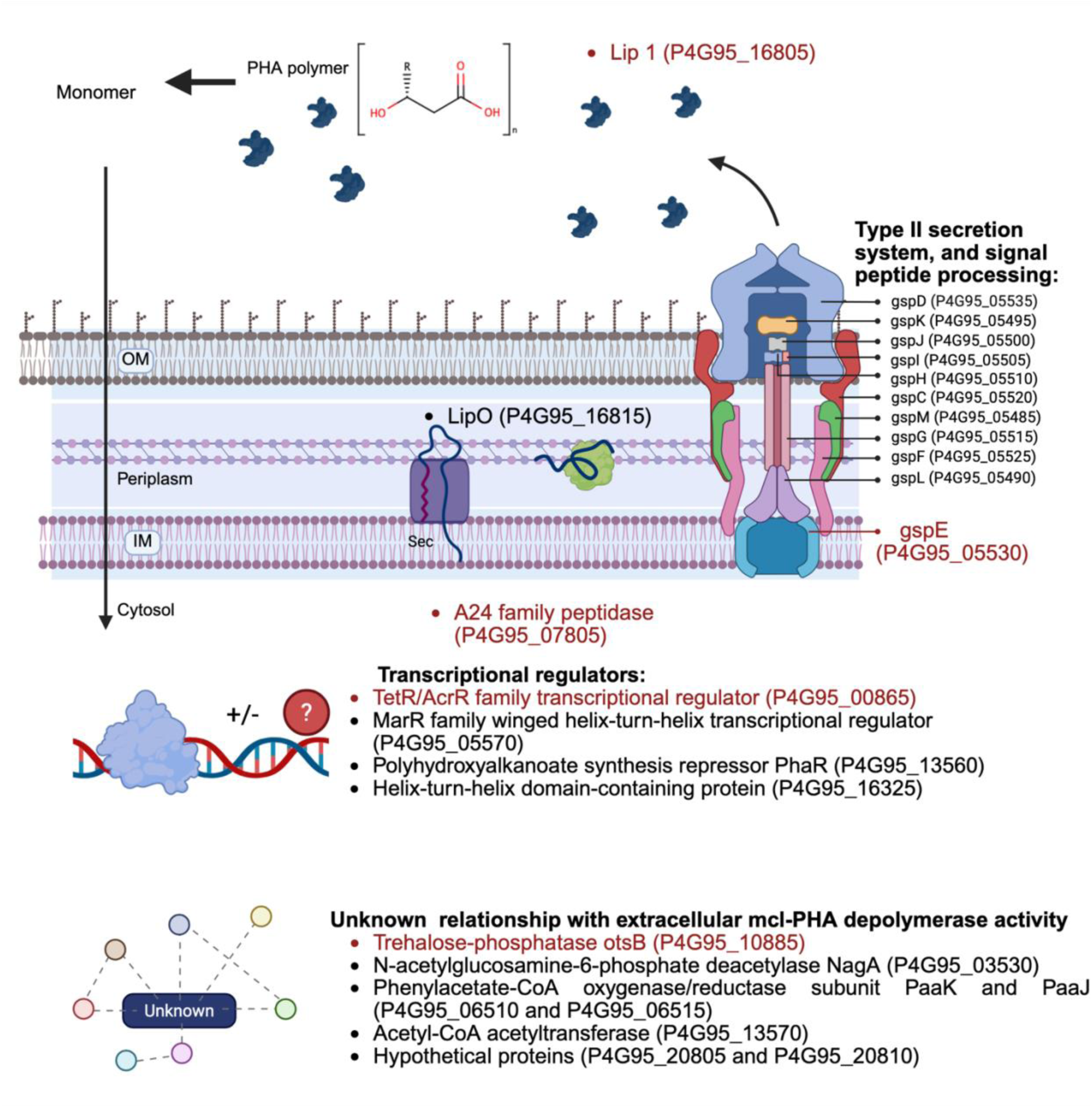
Proposed model of components involved in *B. vietnamiensis* LMG 16232 extracellular mcl-PHA degradation activity. The elements in this model were deduced from the transconjugants that either lost or showed severely reduced extracellular mcl-PHA depolymerase activity due to transposon insertion. The components where the text is in red have been verified via the creation of insertional mutagenesis using RhaCAST. The regulatory elements influencing this activity are TetR/AcrR family transcriptional regulator (P4G95_00865), Polyhydroxyalkanoate synthesis repressor (P4G95_13560) and HTH-type regulators (P4G95_05570 and P4G95_16325). The Sec system is responsible for transporting the unfolded protein into the periplasmic, as the protein is predicted to contain a sec tag. The putative lipase chaperone (P4G95_16815) is responsible for folding the putative lipase into a mature protein and excreted to the outside of the cell via the T2SS. *B. vietnamiensis* LMG 16232 extracellular mcl-PHA degradation activity is also affected by the transposon interruption in genetic elements related to polyhydroxyalkanoate production and several genes involved in metabolism.

One of the putative regulatory elements identified in this work is a TetR/AcrR family transcriptional regulator CDS (P4G95_00865). This type of regulator is widely distributed among bacteria and controls various cellular processes, including the metabolism of lipid compounds, such as the synthesis and degradation of fatty acid and the storage of polymer (45). The involvement of other regulatory elements, such as HTH-type transcriptional activators and a PHA synthesis repressor, suggests that extracellular mcl-PHA degradation activity of *B. vietnamiensis* LMG 16232 may be integrated with other metabolic pathways. While we did not identify a specific genetic element directly responsible for the uptake of mcl-PHA degradation products, we propose passive diffusion. This hypothesis is based on the structural similarity between mcl-PHA monomers and fatty acids, which both contain a hydrophobic alkyl chain (46). Medium-chain fatty acids (C_7_-C_11_) can enter bacteria through simple diffusion without the involvement of active transport systems or an unknown mechanism (47, 48).

Enzymes that degrade extracellular mcl-PHA and lipases exhibit similar characteristics, including the presence of the lipase box and a conserved non-catalytic amino acid triad (16). Consequently, this study suggests that the mcl-PHA degrading enzyme from *Burkholderia* are lipases that may also possess the capability to degrade mcl-PHA. However, Lip2 (P4G95_16810) is annotated as a lipase, contains the lipase box and is highly similar to Lip1, yet our experimental evidence indicates this lipase cannot degrade mcl-PHA. Our structural analysis of Lip1 and Lip2, based on AlphaFold predictions, highlights a conserved core architecture and variable N- and C-terminal regions, suggesting potential regulatory or accessory roles. Superposition with the *B. cepacia* lipase PDB structure 4LIP demonstrated overall structural similarity but also revealed an obstructive alpha-helix in Lip1 and Lip2, which is absent in 4LIP, and may influence enzyme activity. While both Lip1 and Lip2 maintain a conserved catalytic triad, consistent with lipase activity, docking experiments with ligands tributyrin and octanoate showed strong and specific binding for Lip1. In contrast, Lip2 exhibited multiple binding modes, indicating a more flexible binding pocket. A difference in the amino acid at position 73: (Lip1 contains glutamine, while Lip2 has tyrosine) is suggestive of differential catalytic activity.

In summary, Lip1 appears to function as an active lipase and mcl-PHA depolymerase, while Lip2 may have altered activity due to structural differences. Additional biochemical and structural studies will be needed to validate these conclusions. Overall, this study demonstrates the power of visual screens combined with high-density transposon mutagenesis to establish gene-to-function links and provides new insight onto the degradation of extracellular mcl-PHA by the genus *Burkholderia*.

## Material and methods

### Strains, Selective Antibiotics, and Growth Conditions

Strains and plasmids are shown in **Table 3**. Primers used in this study are listed in **Table 4**. All strains were grown in LB-Lennox medium (Difco). Strains of *Burkholderia* and the strains of *E. coli* were initially grown at 37 °C, except for *E. coli* Stbl2, which was grown at 30 °C. The following selective antibiotics were used: chloramphenicol (Sigma; 20 μg/mL for *E. coli*), trimethoprim (Sigma; 100 μg/mL for strains of *Burkholderia*, 50 μg/mL for *E. coli*), gentamicin (Sigma; 50 μg/mL for *Burkholderia*). *Genomic and plasmid extraction* Genomic DNA of the *B. vietnamiensis* LMG 16232 mutants made with RhaCAST were extracted using the PureLink microbiome DNA purification kit (Invitrogen). Genomic DNA of the Tn-seq pools were extracted using standard isopropanol precipitation (49). Plasmid DNA was extracted using the EZNA Plasmid DNA Mini Kit (Omega Bio-tek). Both genomic and plasmid DNA were eluted or solubilized in Tris-HCl pH 8.5.

**Table 4.**
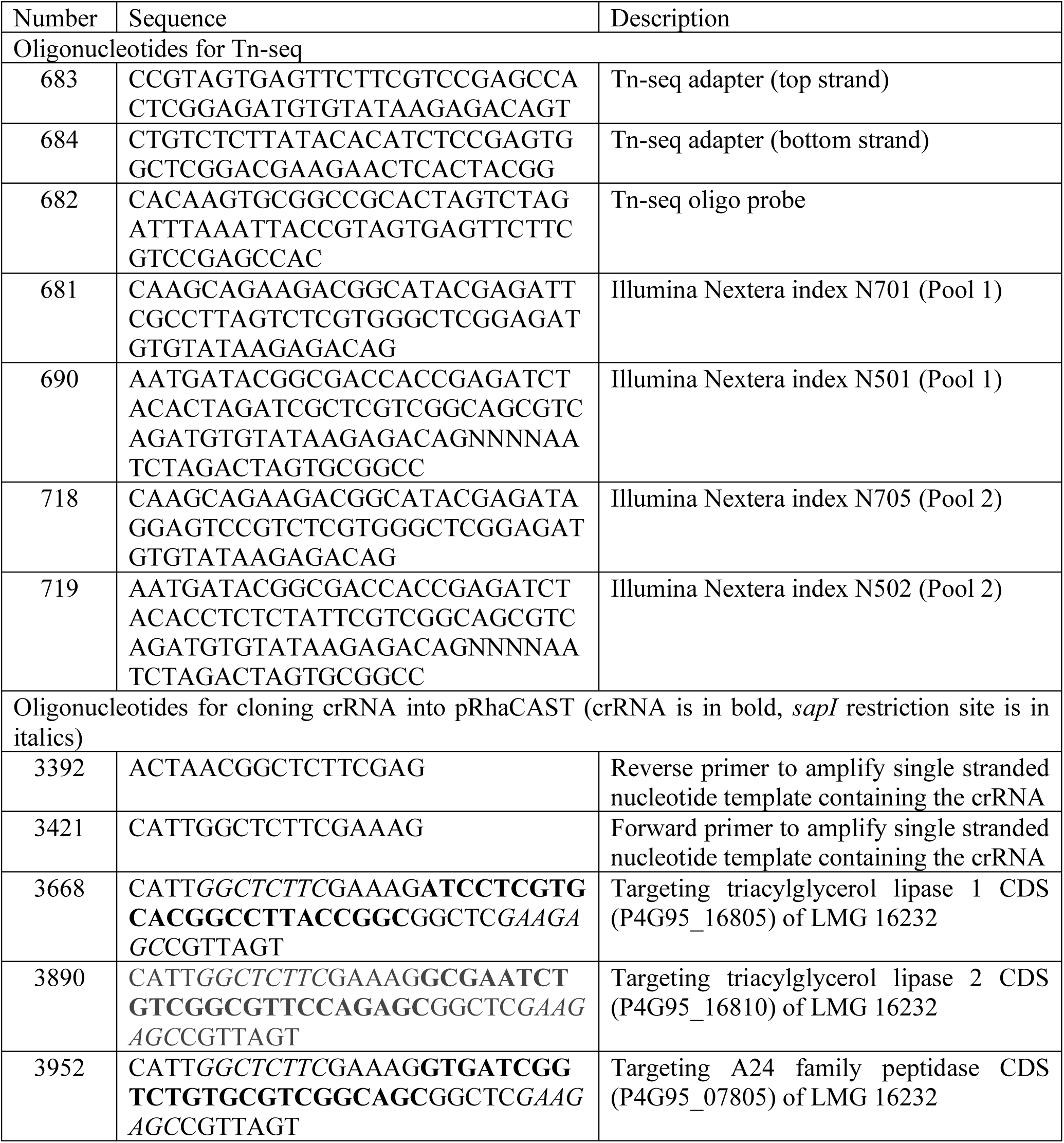
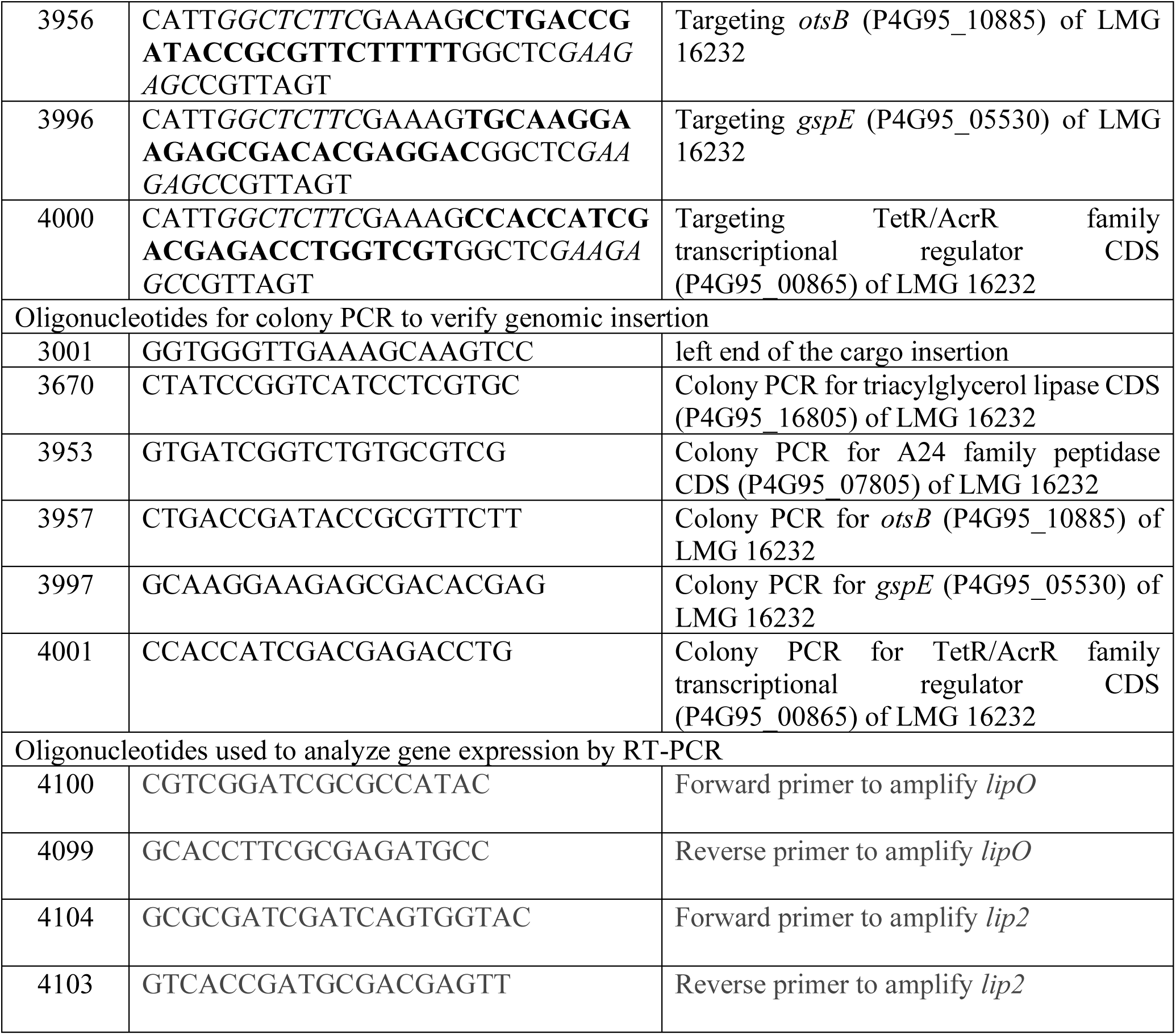
Primers/templates used in this study.

### Substrate and media preparation

The poly(3-hydroxyoctanoate) (PHO) (Product name: VersaMer-8-90) colloidal suspension was created by dissolving 400 mg of PHO (Polyferm Canada Inc., Canada) into 200 mL acetone. Once the PHO was dissolved, 40 mL of dH_2_O was added. The acetone was evaporated off using a speed evaporator. The colloidal suspension was stored at 4° C until needed.

The mcl-PHA containing medium consisted of an M9 minimal medium agar (BD Difco, USA) with 2 mM phenylalanine, 2 mM MgSO_4_, 10 mg/ mL of PHO and 0.3 µg/mL Nile red. PHO was added to the medium by using a previously prepared colloidal suspension of PHO. The PHO, Nile red, MgSO_4_, and phenylalanine were added after autoclaving to melted M9 agar. 25 mL of medium was then poured into petri plates. After solidifying, the plates were stored at 4°C until needed.

Tributyrin medium was made using a minimal medium with 2 mM phenylalanine and 2 mM MgSO_4_ with tributyrin added to a final concentration of 4% (v/v) to the medium according to the Sigma-Aldrich instructions (Sigma, St. Louis, Missouri, USA). The tributyrin was added after autoclaving whilst the agar medium was around 80°C. The medium with tributyrin was vigorously mixed until ready to pour into petri plates. After solidifying, the plates were stored at 4°C until needed.

### Screening for mcl-PHA and tributyrin degradation

To determine the strain’s ability to degrade mcl-PHA and tributyrin degradation. The overnight cultures were grown at 30°C and washed twice with PBS. 20 µL of the culture was inoculated after being adjusted to an OD_600_ of 1. For mcl-PHA degradation determination, the plates were incubated for 7-20 days; For tributyrin degradation determination, the plates were incubated for 3 days, both incubated at 30°C. The visualization of the halo degradation was performed using AlphaImager EP (Cell Biosciences, Santa Clara, CA). For the mcl-PHA plates containing Nile red, the image was taken under UV excitation. Colonies that appeared to have lost the PHO degradation activity on the first round of screening were then subjected to a secondary screening to rule out false positives. 66 and 18 transconjugants were isolated without or with weak PHO degradation activity, respectively.

### Transposon mutagenesis, Tn-seq library preparation and genome mapping

*B. vietnamiensis* LMG 16232 was subjected to transposon mutagenesis with pRBrhaBoutgfp (50). A total of 50,000 single colonies were screened. This number is based on the expected average transposon insertion density from the model, approximately eight transposon insertions per gene. Assuming a genome with N = 6000 non-essential genes and a collection of N = 50,000 transposon insertion mutants, the probability that a single gene will see no transposon insertions is: P(0, k, N) = e^(-50,000/6,000) = 2.4×10^-4^; the total number of genes that will see no insertions is: N* e^(-k/N) = 6,000 x 2.4×10^-4^ = 1.4 genes; and the total number of genes will be found with this transposon density is n(insertion) 6,000* (1-e ^(-50,000/ 6,000)) = 5,999 genes.

Tn-seq was chosen to identify the transposon insertion sites. To enrich transposon-genome junctions and to determine the insertion site, an adapted Tn-seq method, Tn-seq circle, developed by Gallagher et al. (51), was performed as previously described with some minor modifications (49, 51). Briefly, cultures of individual mutants were grown individually to approximately OD_600_ of 3 and were combined into two separate pools, based on without (Pool 1) or with weak (Pool 2) PHO degradation activity. The cultures were harvested, followed by the genomic DNA extraction. The DNA was sheared to an average size of 300 bp via ultrasonication with a Covaris M220 (duty factor of 20 %, a peak incident power of 50W, 200 cycles per burst, and a duration of 75s), followed by end repair with the NEBNext End Repair Module (NEB) and A-tailing. Next, the products were ligated with adaptors, consisting of annealing oligos 683 and 684 with the Quick Ligation Kit (NEB) and digested with *PacI* (NEB). Probe 682 in the presence of Ampligase (Lucigen) was used to circularize fragments containing the transposon sequence. The exonuclease treatment was carried out with a mix of T7 gene 6 exonuclease (ThermoFisher), Lambda exonuclease (NEB), and Exonuclease I (NEB) to digest the linear fragments. The enrichment of the transposon–genome junctions was confirmed by comparing the amplification of the transconjugant’s pools with wild-type *B. vietnamiensis* LMG 16232 using qPCR with iTaq Universal SYBR Green Supermix (Bio-Rad). The number of PCR cycles corresponding to ∼50% saturation by qPCR was chosen to amplify transposon-genome junctions with iTaq DNA polymerases (Bio-Rad) for Illumina sequencing. PCR primers 681, 690, 718, and 719 contain the Nextera indices and were used in appropriate pairs (**Table 4**). The indexed samples were then cleaned up with SeraMag beads (Cytiva), analysed on a TapeStation4150 (Agilent Technologies) and sequenced with an Illumina MiSeq 2500. Raw reads were deposited in the NCBI Sequence Read Archive (SRA) repository and will be publicly available after publication under BioProject: PRJNA1082765, accession: SRR28216496 (Pool 1), SRR28216495 (Pool 2).

The sequenced reads of these mutants were mapped to the *B. vietnamiensis* LMG 16232 genome with the Galaxy bioinformatics platform (https://usegalaxy.org). Using Cutadapt (Galaxy v2.9+galaxy0) (52), the transposon sequence 5’-NNNNAATCTAGACTAGTGCGGCCGCACTTGTGTATAAGAGTCA-3’ was removed from the FASTQ file. Next, using Burrows-Wheeler transform (BWA) (Galaxy v0.7.17.4) (53), the reads were mapped to the close genome of *B. vietnamiensis* LMG 16232 (29). The BWA’s BAM output was then converted to SAM using BAM-to-SAM (Galaxy 2.0.2) (54). The output was visualized using Geneious 8.1.9.

### Determination of the PHO content beneath inoculated culture

After 4 days, the bacterial growth from the inoculated culture from the phenotypic assay was scraped off from the agar. An agar disk beneath the inoculated culture was cut using the back of the P1000 pipette tip. For control, a disk was cut where no cell was inoculated. The samples were dried at 60°C for 16 hours, and the PHO was extracted in chloroform using the acid-catalyzed methanolysis procedure as previously described (55, 56). The determination of PHO in the agar was performed using an Agilent 7890A gas chromatograph equipped with a DB-23 capillary column (30 m × 250 μm × 0.25 μm) and a flame ionization detector (Agilent Technologies, Santa Clara, CA). Method operating parameters and peak quantification have been previously described (55, 57). The retention times and response factors for methyl 3-hydroxyoctanic acid were confirmed using standards purchased from Cayman Chemical (Catalogue # 24609).

### Gene expression analysis by reverse transcriptase (RT)PCR

To determine the effect of RhaCAST insertion in *lip1* and *lip2* on the expression of downstream genes, RT-PCR was performed. Briefly, *B. vietnamiensi*s LMG 16232 Wildtype, *lip1*, and *lip2* mutants were grown overnight on LB media at 30°C, the cultures were washed twice with PBS and adjusted to OD_600_ of 1 (Time 0). Twenty µL of the adjusted cultures were then plated onto PHO plates. Bacterial growth was harvested from the plates at days 4 and 7 and the RNA was extracted following PureLink RNA Mini Kit protocol (Invitrogen^TM^) following the manufacture’s instructions for bacterial cells. The RNA was then converted to cDNA using a SuperScript IV VILO Master Mix Kit (ThermoFisher). The PCR reactions were performed with the Q5 High-Fidelity DNA polymerase (New England Biolabs) following the manufacturer’s instructions.

### In silico analysis

Signal peptides were predicted for predicted PHA depolymerases using Signal P 5.0 (https://services.healthtech.dtu.dk/service.php?SignalP) with default settings using both Gram-negative and Gram-positive predictions (58). The alignment of amino acid sequences was performed with Clustal Omega from Geneious Prime®2026.0.2 using default settings. Protein conformation was predicted using ColabFold 2 (59). Molecular docking studies were conducted using Autodock 4 (60) to investigate the interaction between the enzyme and its potential substrates. The docking protocol was guided by the crystal structure of the reference protein (PDB ID: 4LIP). Visual representations of the predicted protein structures and docking interactions were generated using PyMOL version 3.0.3 (61).

### Construction of B. vietnamiensis LMG 16232 insertional mutants

Genes were interrupted using the RhaCAST method reported previously (31). Briefly, 24 bp crRNAs were designed from the genes to be targeted, and the templates were ordered as single-stranded oligos and PCR-amplified using PCR primers 3392 and 3421 to create double-stranded DNA fragments. The PCR products were then introduced into pRhaCAST using Golden Gate cloning with SapI (NEB). The newly constructed plasmids were then transformed into *E. coli* Stbl2. pRhaCAST was then introduced to *B. vietnamiensis* LMG 16232 by tetra-parental mating (31). *E. coli* MM290/pRK2013 helper strain, *E. coli* Stbl2/pRhaCAST with corresponding crRNA, and *E. coli* PIR1/pDonorTp was conjugated into *B. vietnamiensis* LMG 16232 on LB agar plate containing 0.05% rhamnose. The plate was incubated at 37°C for 24 hours. Successful exconjugants were selected on LB plates containing trimethoprim and gentamycin to select against the *E. coli*. Colonies that appeared within 48 h at 37°C were screened by colony PCR. All insertional mutants constructed with RhaCAST were verified by whole genome sequencing using Oxford Nanopore Technologies platforms with v14 library prep chemistry and R10.4.1 flow cells by Plasmidsaurus (Oregon, USA). The genomic sequence of the insertional mutants is available after under BioProject: PRJNA1205771.

## Author Contributions

STC and ZLY formulated the ideas and design of the project; and EQ performed GGL performed docking experiment; ZLY, SY and MS designed and created plasmids and mutants; ZLY, SY, MS, RD, and WB performed phenotypic assays; ZLY and AMH performed library prep for Tn-seq. EQ performed gene expression analysis. ZLY and AM processed and analyzed the Tn-seq data; ZLY and STC wrote the manuscript and edited together with SY, EQ, GGL, MS, RD, AMH, AM, WB, RS, DBL, and DFDP; STC, DFDP, DBL, RS, and WB provided financial support and supervised the work.

## Acknowledgements and Funding Sources

This work was supported by a Natural Sciences and Engineering Research Council (NSERC) of Canada Discovery Grant to SC and a University of Manitoba Collaborative Research Program (UCRP) grant to SC and DBL. ZLY was supported by a Research Manitoba and University of Manitoba Graduate Fellowship (UMGF); MS and SY were supported by NSERC Undergraduate Student Research Awards (USRA); AMH was supported by a Cystic Fibrosis Canada studentship and a Canadian Institutes of Health Research (CIHR) Vanier award.

