## Supplementary material for "*Burkholderia vietnamiensis* Genes Involved in Extracellular Medium-Chain-Length Polyhydroxyalkanoate Degradation": Suplemental material

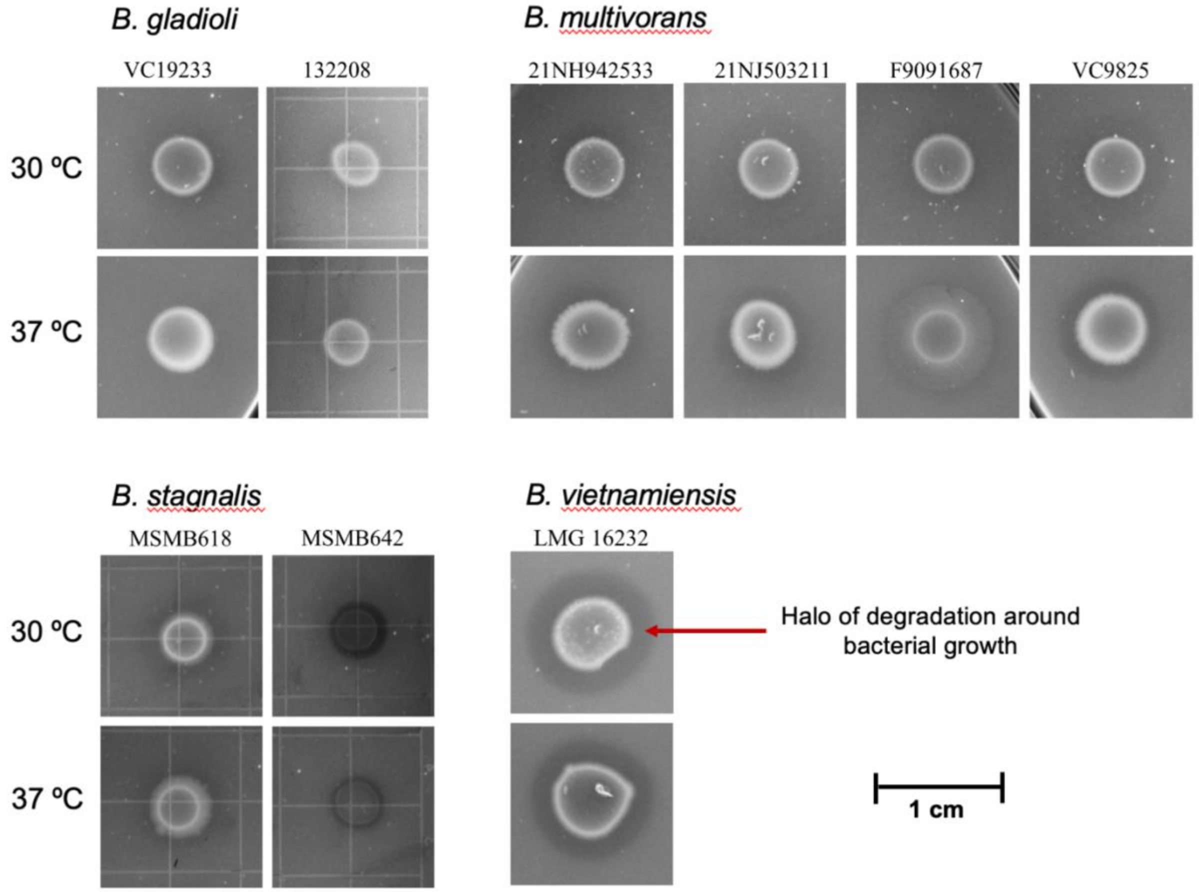


**Fig S1.** The mcl-PHA degradation activity of *Burkholderia* strains. The strains were grown on agar plates containing Nile red and PHO for 20 days at 30°C and 37°C. The phenotypes were visualized under UV irradiation, and the mcl-PHA degradation activity was ranked based on the following scale: (–) no clearing, (+) weak clearing, (++) moderate clearing. The phenotypes observed for each strain were as follows: *B. gladioli* VC19233 showed a rating of (+) at both temperatures, while *B. gladioli* 132208 exhibited (+) at 30°C and (-) at 37°C. Both *B. multivorans* strains, 21NH942533 and VC9825 displayed a rating of (+) at both temperatures, while *B. multivorans* 21NJ503211 and showed (+) at 30°C and (-) at 37°C. Finally, *B. stagnalis* MSMB618, *B. stagnalis* MSMB642, and *B. vietnamiensis* LMG 16232 were rated (++) at both temperatures. The arrow indicates the halo of degradation. A scale bar (1 cm) is shown for reference.

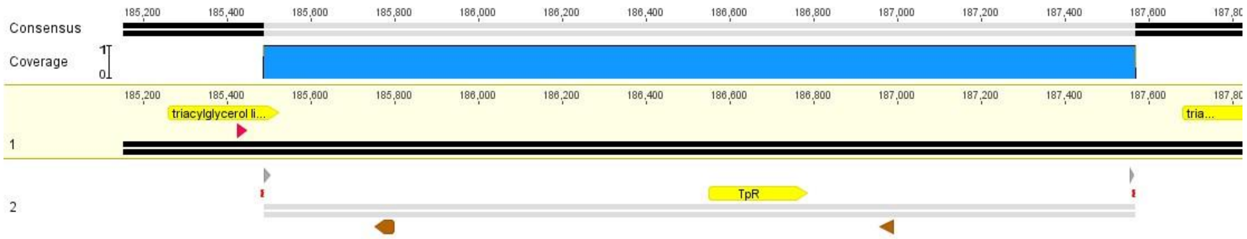


**Fig S2.** Alignment of nanopore sequencing of *B. vietnamiensis* LMG 16232 transconjugants with genomic integration in triacylglycerol lipase CDS (locus tag: P4G95_16805) created using the RhaCAST system. The cargo region flanking the left end (LE) and right end (RE) of the plasmid pDonorTp was mapped and aligned to the sequenced genome. The pink arrows indicate the position of crRNA. Transposon's left end (LE) and right end (RE) elements are represented in a grey arrow. Trimethoprim resistance cassette (TpR) was situated between FRT recognition sites (brown arrow). Mutant created with crRNA 3668, distance from PAM: 63 nt.

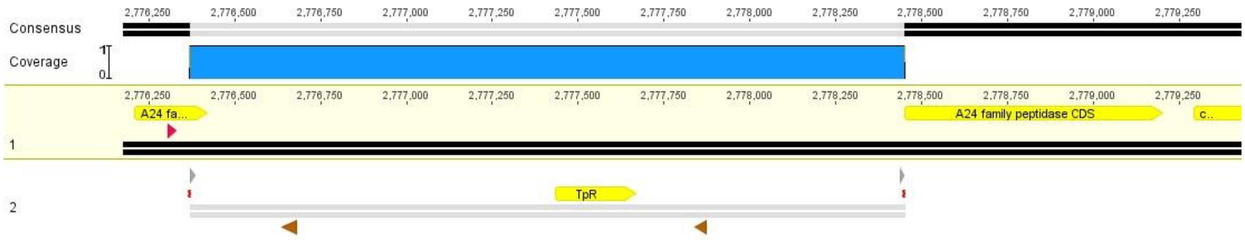


**Fig S3.** Alignment of nanopore sequencing of *B. vietnamiensis* LMG 16232 transconjugants with genomic integration in A24 family peptidase CDS (locus tag: P4G95_07805) created using the RhaCAST system. The cargo region flanking the left end (LE) and right end (RE) of the plasmid pDonorTp was mapped and aligned to the sequenced genome. The pink arrows indicate the position of crRNA. Transposon's left end (LE) and right end (RE) elements are represented in a grey arrow. Trimethoprim resistance cassette (TpR) was situated between FRT recognition sites (brown arrow). Mutant created with crRNA 3952, distance from PAM: 64 nt.

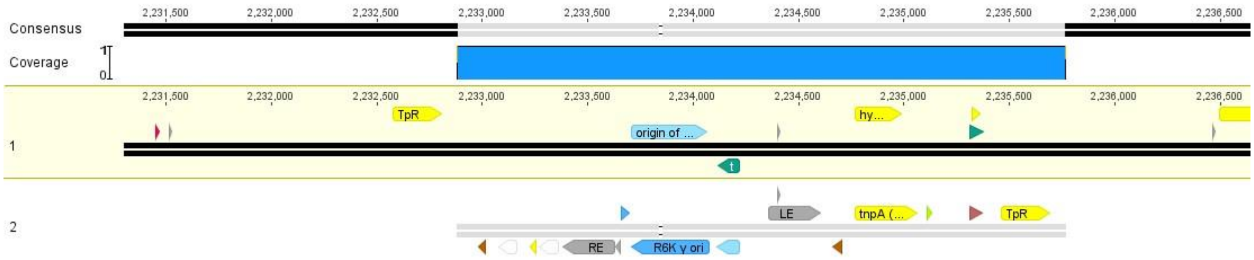


**Fig S4.** Alignment of nanopore sequencing of *B. vietnamiensis* LMG 16232 transconjugants with genomic integration in *otsB* (locus tag: P4G95_10885) created using the RhaCAST system. The pink arrows indicate the position of crRNA. Transposon's left end (LE) and right end (RE) elements are represented in a grey arrow. Trimethoprim resistance cassette (TpR) was situated between FRT recognition sites (brown arrow). This mutant has co-integration of the whole pDonorTp plasmid. Mutant created with crRNA 3956 distance from PAM: 65 nt.

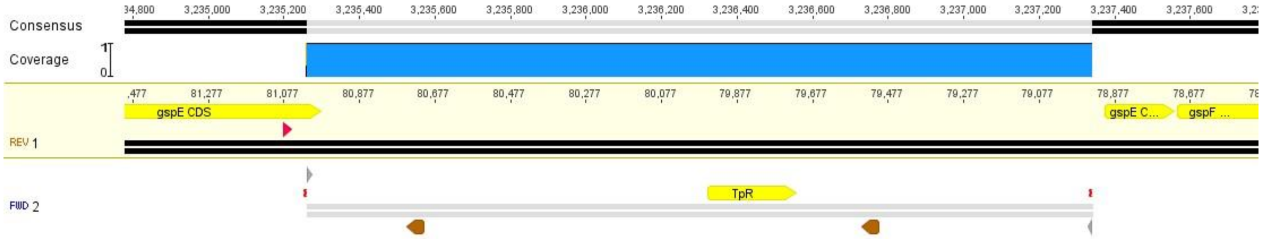


**Fig S5.** Alignment of nanopore sequencing of *B. vietnamiensis* LMG 16232 transconjugants with genomic integration in *gspE* (locus tag: P4G95_05530) created using the RhaCAST system. The cargo region flanking the left end (LE) and right end (RE) of the plasmid pDonorTp was mapped and aligned to the sequenced genome. The pink arrows indicate the position of crRNA. Transposon's left end (LE) and right end (RE) elements are represented in a grey arrow. Trimethoprim resistance cassette (TpR) was situated between FRT recognition sites (brown arrow). Mutant created with crRNA 3996, distance from PAM: 63 nt.

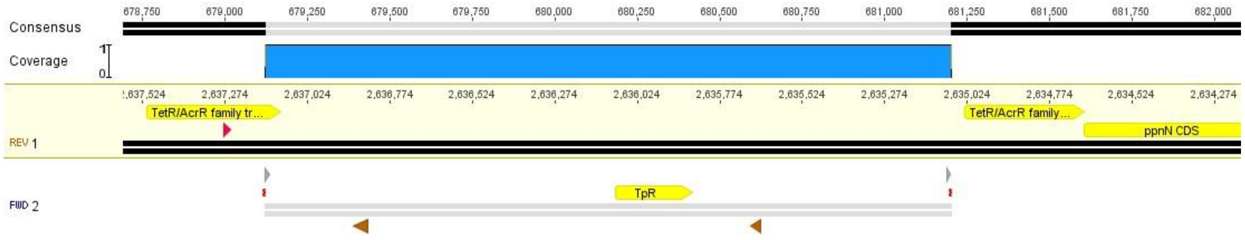


**Fig S6.** Alignment of nanopore sequencing of *B. vietnamiensis* LMG 16232 transconjugants with genomic integration in TetR/AcrR family transcriptional regulator CDS (locus tag: P4G95_00865) created using the RhaCAST system. The cargo region flanking the left end (LE) and right end (RE) of the plasmid pDonorTp was mapped and aligned to the sequenced genome. The pink arrows indicate the position of crRNA. Transposon's left end (LE) and right end (RE) elements are represented in a grey arrow. Trimethoprim resistance cassette (TpR) was situated between FRT recognition sites (brown arrow). Mutant created with crRNA 4000, distance from PAM: 126 nt.

```
P4G95_16805      1 MVRSMRSRVVAAAVACAMSGAQFAGTTAVMTLATTHAATAATAATDDYAATRYPVILVHG   60
                   M +SMRSRV+A A ACAMS A FAG TAVM LATT  A AATA +D YAATRYP++ VHG
PG95_16810       1 MTKSMRSRVMAGAAACAMSAAPFAGATAVMALATTRTAVAATAPSDTYAATRYPLVFVHG   60

P4G95_16805     61 LTGTDKYAGVLEYWYGIQEDLQRHGATVYVANLSGFQSDDGPNGRGEQLLAYVKTVLAAT  120
                   + G +K  G ++ WYGI+ DLQ+HGA+VYVANLS FQSDDGPNGRGEQLLAYVK VLAA+
PG95_16810      61 INGNEKIGGAIDQWYGIRGDLQQHGASVYVANLSAFQSDDGPNGRGEQLLAYVKRVLAAS  120

P4G95_16805    121 GATKVNLIGHSQGGLTSRYVAAVAPDLVASVTTIGTPHRGSEFADFVQSVLAYDPTGLSS  180
                   GAT+VNL+GHSQGGLTSRYVAAVAP+LVASVTTIGTPHRGSEFADFVQ VLAYDPTGLSS
PG95_16810     121 GATRVNLVGHSQGGLTSRYVAAVAPELVASVTTIGTPHRGSEFADFVQRVLAYDPTGLSS  180

P4G95_16805    181 SVIATFFNVFGILTSSSNNTNQDALAALKTLTTSQAAAYNQNYPSAGLGAPGSCQTGAPT  240
                   +VIA F N+   LTS++ N  QDALAAL+TLTTSQAA YN NYPSAGLGAPGSC+TGA T
PG95_16810     181 AVIAKFVNLMAFLTSTNFNLKQDALAALQTLTTSQAAVYNVNYPSAGLGAPGSCETGAST  240

P4G95_16805    241 ETVGGNTHLLYSWAGTAIQPTVSMFGVTGAKDVSTIPVIDPANALDLSTLALLGTGTVMI  300
                   ET+GG+THLLYSW GTAI+P  S+ GV  AKDVSTIP+IDPA   D STLA+L TGTVMI
PG95_16810     241 ETIGGHTHLLYSWTGTAIRPAFSLSGVMLAKDVSTIPLIDPAYKDDSSTLAMLATGTVMI  300

P4G95_16805    301 NRGAGQNDGLVSKCSALYGQVLGTRYKWNHLDEINQLLGVRGAYAEDPVAVIRTHANRLK  360
                   NRG+G+NDGLVSKCSALYGQVLGT YKWNH+DEINQ LGV GA+AEDPVAVIRTHANRLK
PG95_16810     301 NRGSGRNDGLVSKCSALYGQVLGTHYKWNHVDEINQTLGVHGAFAEDPVAVIRTHANRLK  360

P4G95_16805    361 LAGV 364
                   LAGV
PG95_16810     361 LAGV 364
```

**Fig. S7**. Amino acid sequence alignment of Lip1(P4G95_168050 and Lip2 (P4G95_16810) of *B. vietnamiensis* LMG 16232. Consensus is shown with red letters; + indicates amino acid similarity. Differences at position 73 (as per Fig. 7F) are noted in blue for tyrosine (Y) in Lip1 and glutamine (Q) in Lip2. The lipid box is bolded. The catalytic triad amino acids S131, N308 and H330 are noted in green. The alignment was generated with Geneious Prime®2026.0.2 (Geneious Alignment) using default settings. Identity = 284/364 (78%); Similarity = 316/364 (86%); Gaps = 0/364 (0%).

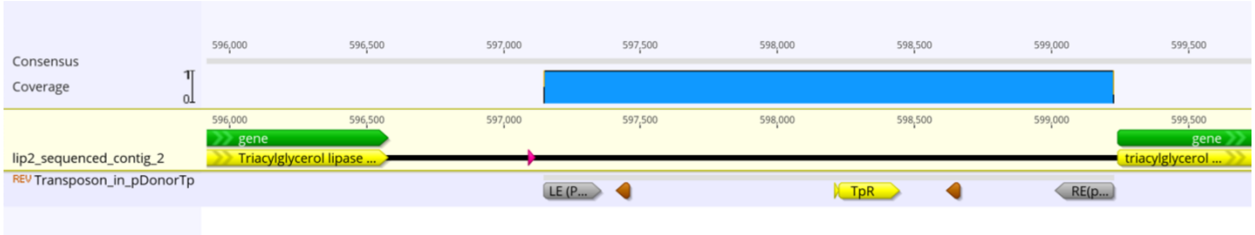


**Fig S8.** Alignment of nanopore sequencing of *B. vietnamiensis* LMG 16232 transconjugants with genomic integration in triacylglycerol lipase CDS (*lip2* locus tag: P4G95_16810) created using the RhaCAST system. The cargo region flanking the left end (LE) and right end (RE) of the plasmid pDonorTp was mapped and aligned to the sequenced genome. The pink arrows indicate the position of crRNA. Transposon's left end (LE) and right end (RE) elements are represented in a grey arrow. Trimethoprim resistance cassette (TpR) was situated between FRT recognition sites (brown arrow). Mutant created with crRNA 3890, distance from PAM: 66 nt.

```
                       10        20        30        40        50        60        70
              ----:----|----:----|----:----|----:----|----:----|----:----|----:----|
Consensus     MXRSMRSRVV---------------AGAXACAMSXA----------PFAGTTAVMTLATTRXA-A-TAPX   43
PG95_16810    MTKSMRSRVM---------------AGAAACAMSAA----------PFAGATAVMALATTRTAVAATAPS   45
P4G95_16805   MVRSMRSRVV---------------AAAVACAMSGA----------QFAGTTAVMTLATTHAATAATAAT   45
AAA50466      MARTMRSRVV---------------AGAVACAMSIA----------PFAGTTAVMTLATTHAAMAATAPA   45
CAE81078      ----MKKLLAGVFGVVAMSLSAQAAKKASNCEVT----GLVDRMTCPYLEKL-VSGPHLTRHVKY-SLPK   60
AFQ93688      ----------------------------------------------------VHTFYH--------PRA    9
AFQ93689      ----------------------------------------------------VHTFYH--------PRS    9
AAA64538      --MPLRTLLCGLLLAVCLGQHALA---ASRCSERPRTLLRPAEVSCSYQSTWLDSGLVGQRKIIY-QTPL   64

                       80        90       100       110       120       130       140
              ----:----|----:----|----:----|----:----|----:----|----:----|----:----|
Consensus     XGYAATRYPVVJVHGGXG--XXDK-YSSLL--XXWY--GIXXDLQXHGAXVYAANLX--XFQ-----GX-   98
PG95_16810    DTYAATRYPLVFVHGING---NEK-IGGAI--DQWY--GIRGDLQQHGASVYVANLS--AFQ--------   97
P4G95_16805   DDYAATRYPVILVHGLTG---TDK-YAGVL--EYWY--GIQEDLQRHGATVYVANLS--GFQ--------   97
AAA50466      AGYAATRYPIILVHGLSG---TDK-YAGVL--EYWY--GIQEDLQQNGATVYVANLS--GFQ--------   97
CAE81078      GKTPKAGWPTVILYQGSLFPVE-FSRSSLMIAGGYNEIRLIQTLLDSGFAVIAPPAIEGVAWMTNIVGI-  128
AFQ93688      MGASGERHPVVIWGNGTG-AVPGI-YSSLL--RHWA--S-------HGFIVAAANTPTSNFALSMRAGID   66
AFQ93689      MGASGERHPVVIWGNGTG-AVPGI-YSSLL--RHWA--S-------HGIIVAAANTPTSNFAISMRAGID   66
AAA64538      GTPPAGGWPVVLIYQGSFFPLNDFSYHSNLPFGGYYEGKLVQNLLDHGYAVIAPSAPADLFWQTNIPGLA  134

                      150       160       170       180       190       200       210
              ----:----|----:----|----:----|----:----|----:----|----:----|----:----|
Consensus     SDDG----PNGRGEQLLAYVKXVFGAXGATRVNLXGHSQGGLXSRYVAXVAPXXVAAVTTXGXPHRXS-E  163
PG95_16810    SDDG----PNGRGEQLLAYVKRVLAASGATRVNLVGHSQGGLTSRYVAAVAPELVASVTTIGTPHRGS-E  162
P4G95_16805   SDDG----PNGRGEQLLAYVKTVLAATGATKVNLIGHSQGGLTSRYVAAVAPDLVASVTTIGTPHRGS-E  162
AAA50466      SDDG----PNGRGEQLLAYVKTVLAATGATKVNLVGHSQGGLSSRYVAAVAPDLVASVTTIGTPHRGS-E  162
CAE81078      -DYDTSEDFYFVEELLVAMGNGEFGKLNMDRLYATGISSGGYHSSRMAVAFPGVFKALAVHSASYADCGG  197
AFQ93688      VLERR---NADPGSE---Y----FGRVDLARIGSAGHSQGGAAAVNAAVD-ARVDTAVPIQPGPLTDP-D  124
AFQ93689      VLERR---NADPGSE---Y----FGRVDLEHIGSAGHSQGGAAAVNAAID-ARVDTAVPIQPGPLTDP-D  124
AAA64538      QAYELSTDYDFLGNVLAAIASGHFGPLNAQRQYATGISSGGYNTSRMAVSFPGKFRALAVQSGSYATCSG  204

                      220       230       240       250       260       270       280
              ----:----|----:----|----:----|----:----|----:----|----:----|----:----|
Consensus     FADFVQXVLA-YD-PTGLSSXVIAXFVNVX-------------------TLT------YNXNYPSAG---  203
PG95_16810    FADFVQRVLA-YD-PTGLSSAVIAKFVNLMAFLTSTNFNLKQDALAALQTLTTSQAAVYNVNYPSAGLGA  230
P4G95_16805   FADFVQSVLA-YD-PTGLSSSVIATFFNVFGILTSSSNNTNQDALAALKTLTTSQAAAYNQNYPSAGLGA  230
AAA50466      FADFVQDVLA-YD-PTGLSSSVIAAFVNVFGILTSSSHNTNQDALAALQTLTTARAATYNQNYPSAGLGA  230
CAE81078      PMCFVPAQVPENHPPTIFLHGRLDPVVPVR-------------------TMY-----PYHETLKNQG---  240
AFQ93688      L--------------TG-----------------------------------------------------  127
AFQ93689      L--------------TD-----------------------------------------------------  127
AAA64538      PLCVVPDQLPADHPPTLFLHGFVDAVVPWW-------------------SMD-----LYYDRLLHQG---  247

                      290       300       310       320       330       340       350
              ----:----|----:----|----:----|----:----|----:----|----:----|----:----|
Consensus     ---CETGAXTETXGGHT------HLLYSWXGTAIXPXVSXFGVXGAXDXSTXPXJDPAXYXDSSTLAXXX  264
PG95_16810    PGSCETGASTETIGGHT------HLLYSWTGTAIRPAFSLSGVMLAKDVSTIPLIDPAYKDDSSTLAMLA  294
P4G95_16805   PGSCQTGAPTETVGGNT------HLLYSWAGTAIQPTVSMFGVTGAKDVSTIPVIDPANALDLSTLALLG  294
AAA50466      PGSCQTGAPTETVGGNT------HLLYSWAGTAIQPTLSVFGVTGATDTSTLPLVDPANVLDLSTLALFG  294
CAE81078      ---VETEMFVSPWARHEWLEEAPELITNWFINHK------------------------------------  307
AFQ93688      --------------------------------------VPVFYLAGERDLTVWPALVKALYRDSD------  154
AFQ93689      --------------------------------------VPMFYLAGQRDLTVWPALVKALYRDSS------  154
AAA64538      ---IETARYTEPLGGHEWFAASPGKVLAWFNAHP------------------------------------  314

                      360       370       380       390       400       410       420
              ----:----|----:----|----:----|----:----|----:----|----:----|----:----|
Consensus     TGTVMINRGXGXNDGLVSKCSALYGZVLGTGYKWNHLDEINQXLGVXGAFAEDPVAVIRTHANRLKLAGV  334
PG95_16810    TGTVMINRGSGRNDGLVSKCSALYGQVLGTHYKWNHVDEINQTLGVHGAFAEDPVAVIRTHANRLKLAGV  364
P4G95_16805   TGTVMINRGAGQNDGLVSKCSALYGQVLGTRYKWNHLDEINQLLGVRGAYAEDPVAVIRTHANRLKLAGV  364
AAA50466      TGTVMINRGSGQNDGLVSKCSALYGKVLSTSYKWNHLDEINQLLGVRGAYAEDPVAVIRTHANRLKLAGV  364
CAE81078      ----------------------------------------------------------------------  271
AFQ93688      -----------------HLPAVYGEVRGAG----HLSS----IGDGGDFRAPTTA---------------  184
AFQ93689      -----------------HLPAVYGEVRGAG----HLSS----IGDGGDFRAPTT----------------  183
AAA64538      ----------------------------------------------------------------------  278
```

**Fig S9.** Amino acid sequence alignment of the *B. vietnamiensis* LMG 16232 Lip1 (P4G95_16805) and Lip2 (P4G95_16810) with the *B. cepacia* lipase (GenBankAAA50466) and mcl-PHA depolymerase sequences from *Streptomyces roseolus* (GenBank: AFQ93688.1), *Streptomyces venezuelae* (GenBank: AFQ93689.1), *Bdellovibrio bacteriovorus* HD100 (GenBank: CAE81078.1) and *Pseudomonas fluorescens* (GenBank: AAA64538.1). Multiple sequence alignment was generated using Clustal Omega from Geneious Prime®2026.0.2 using default settings. The lipase box, consisting of the pentapeptide sequence (G-X1-S-X2-G), is indicated in red. The signal peptide of *B. vietnamiensis* LMG 16232, predicted by the SignalP (version 5.0) program (58). is indicated in orange. The catalytic triad amino acids serine (in the lipase box), asparagine, and histidine, noted in Fig. 7 as S131, D308 and H220, are indicated in green.

(A)

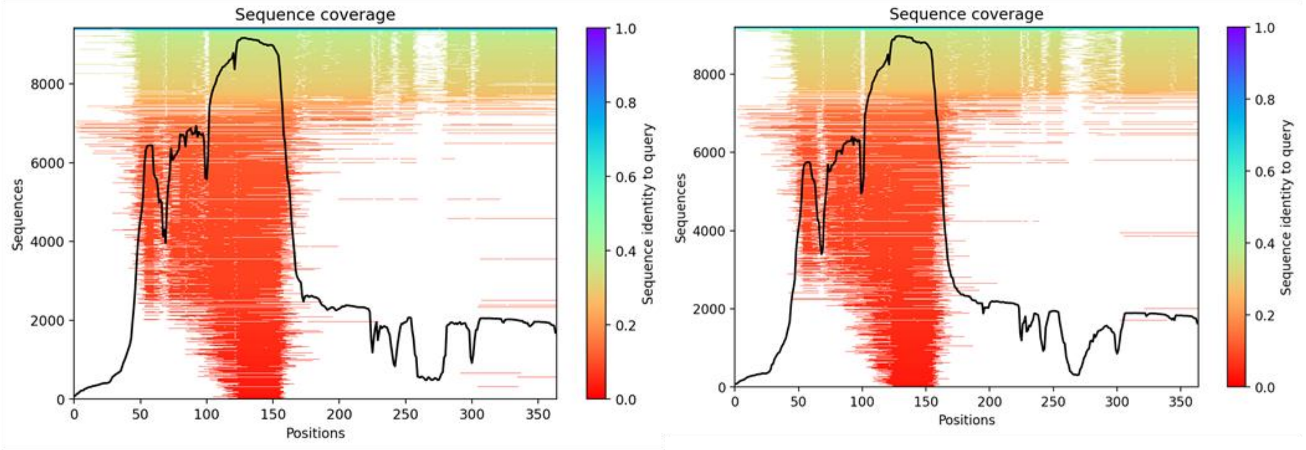
Sequence coverage
Sequences
Positions
Sequence identity to query
Sequence coverage
Sequences
Positions
Sequence identity to query

(B)

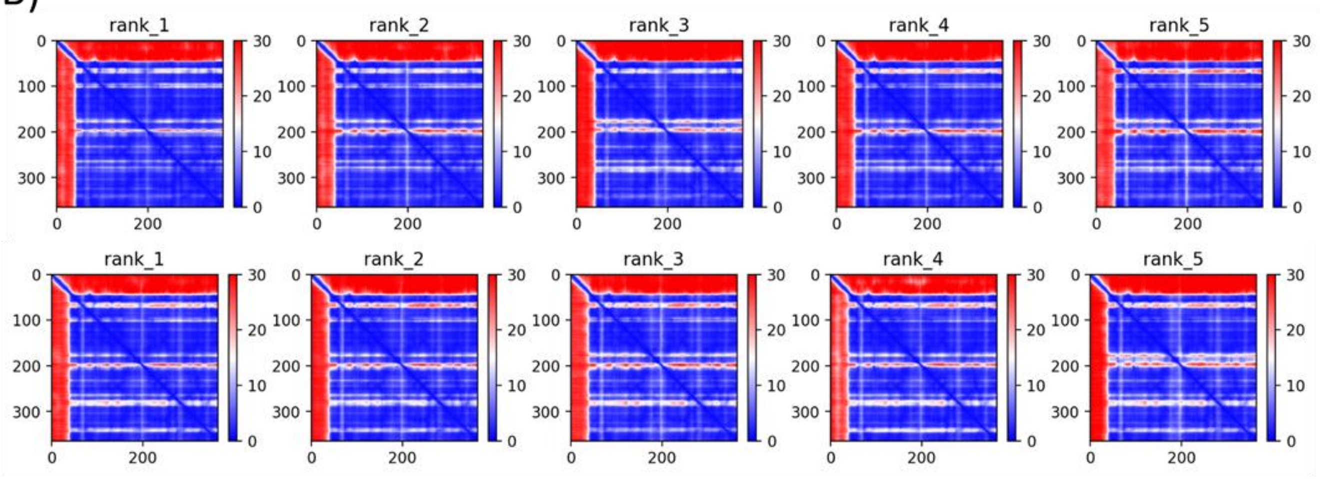
rank_1
rank_2
rank_3
rank_4
rank_5
rank_1
rank_2
rank_3
rank_4
rank_5

(C)

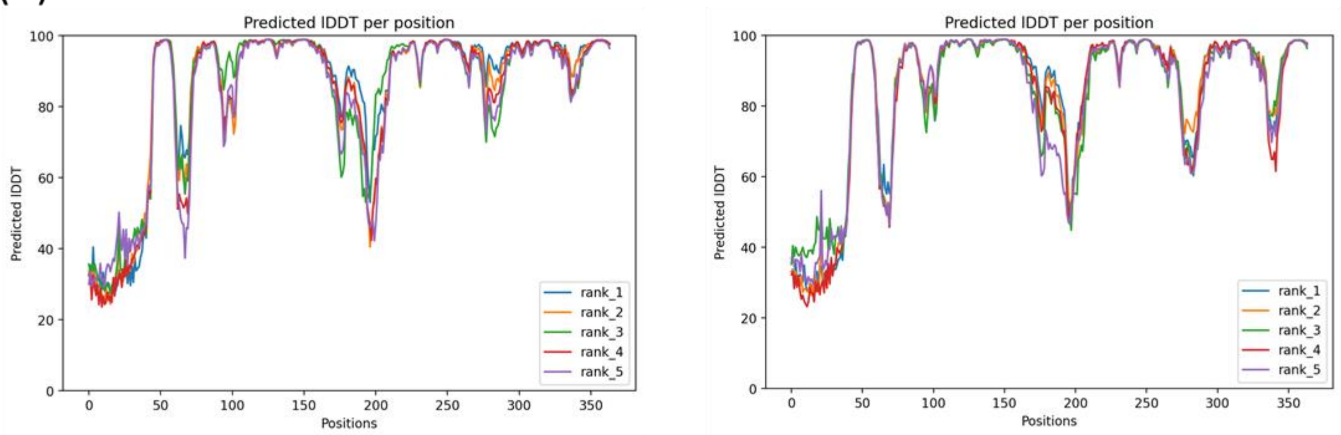
Predicted lDDT per position
Predicted lDDT
Positions
rank_1
rank_2
rank_3
rank_4
rank_5
Predicted lDDT per position
Predicted lDDT
Positions
rank_1
rank_2
rank_3
rank_4
rank_5

**Fig S10.** Sequence conservation, inter-residue error estimates, and structural confidence of Lip1 and Lip2. **A**) Sequence coverage analysis. The black trace represents the number of aligned sequences at each position, while the heatmap colour scale reflects sequence identity to the query. A highly conserved core region (~50–160) is evident, with lower coverage and higher sequence variability observed in the N- and C-terminal regions for both Lip1 (left) and Lip2 (right); **B**) Predicted Aligned Error (PAE) matrices. PAE matrices for the five top-ranked AlphaFold models illustrate inter-residue distance prediction errors for Lip1 (top) and Lip2 (bottom). Low error values (blue) indicate well-defined structural relationships, while higher errors (red) suggest flexible or ambiguous regions. The consistent patterns across models support a stable domain architecture with potential inter-domain flexibility; **C**) Predicted Local Distance Difference Test (pLDDT) scores. The pLDDT values for Lip1 and Lip2 reveal high structural confidence (>90) in the core region, while lower scores in the ~50–100 and ~200–250 regions suggest flexible or intrinsically disordered segments. Trends are consistent across both proteins, reinforcing the reliability of the structural predictions.

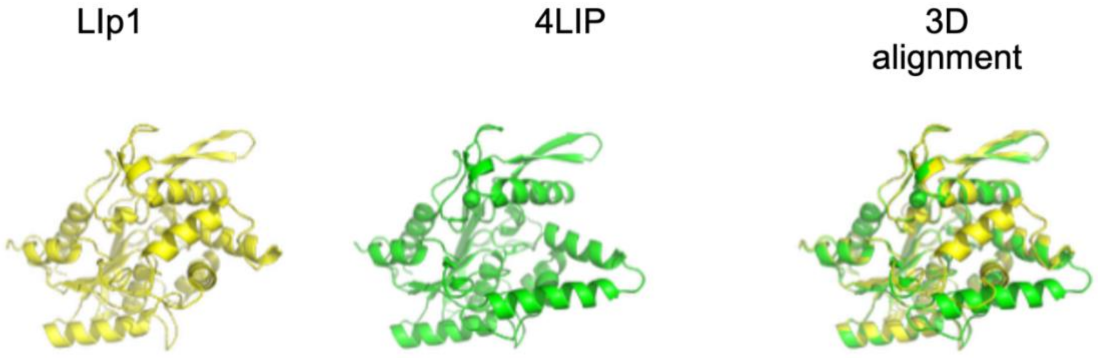


**Fig S11.** Structural comparison between the predicted Lip1 model and the crystallized 4LIP template**.** Superposition of the AlphaFold-predicted structure of Lip1 (yellow) and the 4LIP crystallographic structure (green), highlighting overall structural conservation and key differences. The AlphaFold-predicted Lip1 structure, in which an additional alpha-helix appears to occlude a cavity observed in 4LIP, suggests potential functional implications. The 4LIP crystallographic structure alone shows the original conformation of the enzyme.

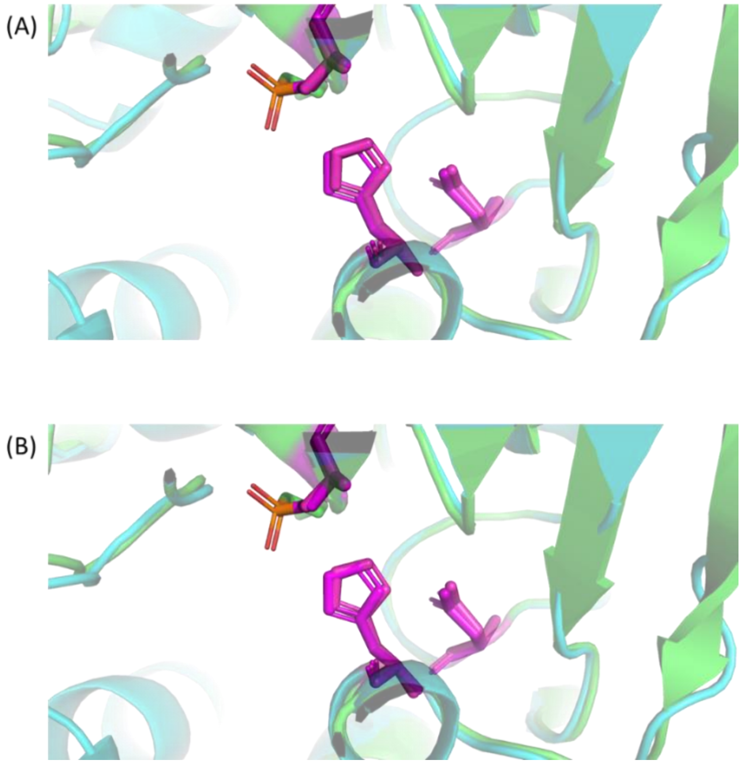


**Fig S12.** Structural superposition of Lip1, Lip2, and 4LIP highlighting the catalytic triad. **A**) Superposition of Lip1 (panel A) and Lip2 (panel B) with the reference lipase 4LIP shows strong conservation of the catalytic triad (highlighted in violet). The preserved spatial arrangement of these key residues suggests that both Lip1 and Lip2 likely retain enzymatic activity.

**Table S1.** Molecular docking results for the evaluated protein-ligand complexes. The table presents the identified binding clusters, along with their lowest and mean binding energies (kcal/mol), and the number of poses (N°) within each cluster (number of runs=100).

| Protein | Ligand | Cluster | Lowest Binding Energy | Mean Binding Energy | N° in Cluster |
|---|---|---|---|---|---|
| Lip1 | OCT | 1 | -4.65 | -4.39 | 100 |
| Lip1 | TRI | 1 | -5.38 | -3.63 | 100 |
| Lip2 | OCT | 1 | -4.68 | -4.38 | 83 |
| Lip2 | OCT | 2 | -3.97 | -4.21 | 12 |
| Lip2 | OCT | 3 | -3.98 | -3.97 | 5 |
| Lip2 | TRI | 1 | -5.32 | -3.65 | 87 |
| Lip2 | TRI | 2 | -4.74 | -4.74 | 1 |
| Lip2 | TRI | 3 | -3.81 | -3.27 | 12 |
